# Astrocytes transform the transcriptomic and epigenetic landscape of developing cortical neurons to promote activity via intrinsic excitability control

**DOI:** 10.64898/2026.05.17.725732

**Authors:** Alison C. Todd, Jing Qiu, Kyle Wardlaw, Kahyee Hor, Sean McKay, Philip Hasel, Xin He, Deepali Vasoya, Mosi Li, Mandy Drake, Owen R Dando, Peter C Kind, Paul S. Baxter, David J. A. Wyllie, Giles E. Hardingham

## Abstract

Astrocytes are implicated in neuronal development and function, particularly of synapses, but their genome-wide impact is unclear. We show that cortical astrocytes trigger epigenetic remodelling in developing cortical neurons and widespread transcriptomic changes distinct from neurons’ intrinsic developmental trajectory. These changes can cause an emergence of neuronal firing activity independent of functional excitatory synaptogenesis. The mechanism is via through astrocyte-derived signals causing the transcriptional repression of inwardly-rectifying K^+^-channels (Kirs), causing a pro-excitatory shift in intrinsic properties. This places neurons in a zone of excitability whereupon cell-autonomous homeostatic control was activated, also associated with Kir regulation. Astrocyte-induced neuronal genes were enriched for schizophrenia, epilepsy, ADHD and Alzheimer’s disease risk, disorders associated with circuit imbalance. Thus, astrocyte-to-neuron signaling can alter firing via intrinsic properties in addition to the classical synaptogenic mechanism. The opposing actions of astrocytes and neuronal activity in regulating neuronal intrinsic properties may prevent hyper- or hypo-activity in neuronal circuits

## Introduction

During CNS development, neurons are first specified from neural stem cells, followed by macroglia (primarily astrocytes and oligodendrocytes). In the mouse forebrain gliogenesis (initially astrocytogenesis) commences at E18, and from then on neurons and astrocytes exist in close proximity to each other. Astrocytes play important tasks in the mature brain to ensure viable and functional neuronal networks. These include providing bioenergetic and redox buffering support ^1–3^ as well as ion homeostasis, pH balance, and uptake of synaptically released neurotransmitters ^4^.

During development there is considerable evidence that neurons and astrocytes influence each other’s maturation. Neurons are responsible for inducing the characteristic stellate morphology of astrocytes in vivo, as well as for inducing the astrocytic expression of glutamate and GABA transporters via signaling pathways including those centred on neuroligin and notch, and neuronal activity can regulate astrocyte metabolism ^5–8^. Considering signaling in the opposite direction (i.e. the influence of astrocytes on neuronal development), the role of astrocytes in controlling excitatory synaptogenesis has been the focus of much interest. Early studies revealed that rat retinal ganglion cells (RGCs) form few functional synapses when cultured on their own, but that synaptogenesis is greatly enhanced when co-cultured with astrocytes ^9,10^. Since then, astrocyte-released factors have been identified as responsible for structural and functional synapse formation, as well as contact-dependent-mechanisms ^7,11–14^.

While astrocyte-dependent synaptogenesis has received a lot of attention, less is known about global influences of astrocytes on developing neurons. For example, it is unclear to what extent astrocytes exert genome-wide effects on the neuronal epigenetic landscape or the transcriptome. While neuronal specification from neural stem cells is well-studied, how astrocytes shape neurons’ developmental trajectory is incompletely understood. Defining non-cell-autonomous control of one cell type’s epigenome or transcriptome by a different cell type is challenging to do since approaches often involve physical separation of cell types which is imperfect and can introduce artefacts due to the effects of cell dissociation and sorting. We previously described a co-culture approach that enables non-cell-autonomous influences to be defined: by establishing co-cultures of distinct cell types from different species (e.g. mouse and rat) then physical cell sorting can be replaced by *in silico* sorting of next-generation sequencing reads ^15^. Here we have applied this approach: maintaining developing cortical rat neurons in the presence or absence of mouse cortical astrocytes in order to define the influence of astrocytes on neuronal epigenetic marks and the neuronal transcriptome, backed up by single-species co-culture single-nucleus RNA-seq. Prompted by analysis of these ‘omic data, uncovered a influence of cortical astrocytes in controlling the excitability of cortical neurons.

## Results

### Astrocytes modify the epigenetic and transcriptomic landscape of developing cortical neurons

To define the genome-wide influence of cortical astrocytes on developing cortical neurons, we used a mixed species co-culture system. E20.5 rat cortical neurons were maintained for 8 days in the presence or absence of mouse astrocytes (for co-cultures, astrocytes were plated down 3 days prior to neuronal platedown) (Fig .1a). The medium conditions were identical (see methods) and in both instances an anti-mitotic agent AraC was included from the point of neuronal plating to prevent any proliferation of rat astrocytes. Neuron-astrocyte co-cultures comprised of approximately 80 % neurons and 20 % astrocytes (Fig. 1a, Fig. S1a,b), whereas neuron mono-cultures comprised of approximately 98 % neurons and <0.5 % astrocytes, measured by immuno-histochemistry and single nucleus RNA-seq cell type profiling (Fig. S1a-e). Both culture preparations were essentially free of microglia (Fig. S1a,b). We seeded an identical volume of the same neuronal cell suspension either onto astrocytes (co-culture) or an empty well (mono-culture). Comparison of neuronal density (Neurochrom positive neurons per field of view) confirms that neuronal density is similar (Fig. S1c)

**Figure 1.**
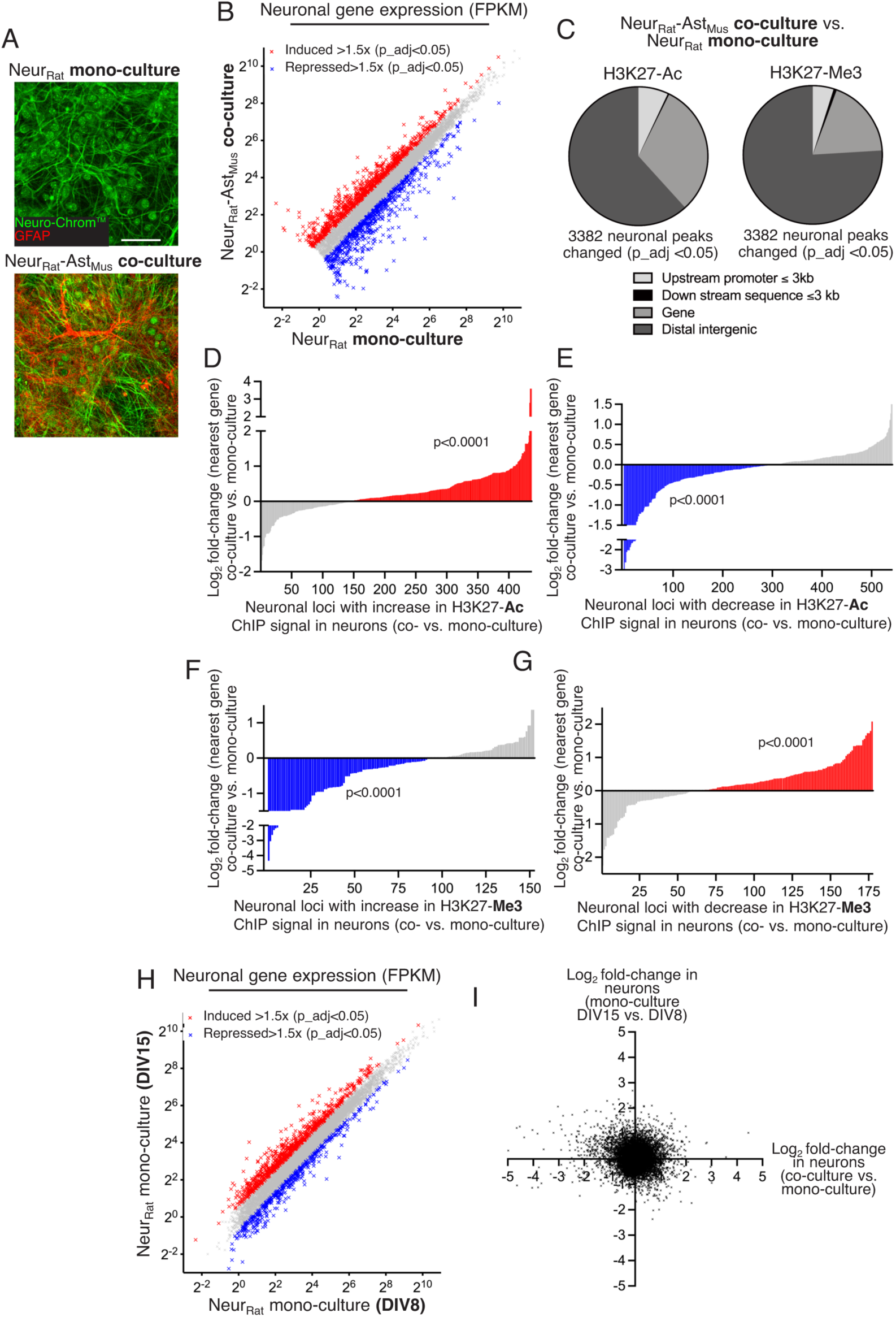
Astrocytes induce changes to the transcriptome of developing cortical neurons. **A)** Example immunofluorescence images of neuronal mono-cultures and neuron-astrocyte co-cultures forming the basis of transcriptomic and epigenomic analyses. Both culture preparations were stained with Neuro-Chrom antibody cocktail (neurons) and anti-GFAP (astrocytes). Scale bar: 50 µm. **B)** RNA-seq was performed on the indicated cultures and both subjected to the same Sargasso workflow to assign reads to rat (neurons). Genes with a DEseq2 fold-change of ≥1.5 fold up (red) or down (blue) due to astrocytes are highlighted (Benjamini Hochberg false discovery rate <0.05, average FPKM ≥1, here and throughout). N=3 per condition. **C)** ChIP-seq was performed on neuronal mono-cultures and neuron-astrocyte co-cultures using the antibodies against H3K27-Ac (left) and H3K27-Me3 (right) and reads separated according to species to define neuronal peaks. The proportions of the peaks relative to genes is shown in the two pie charts **D,E)** H3K27-Ac ChIP peaks significantly increased (D) or decreased (E) in neurons by the presence of astrocytes whose location was in the gene or within 3 kb of the transcription start site were taken and the Log2 fold-change of mRNA expression plotted for genes where average expression was ≥ 1 FPKM. A significant positive astrocyte effect was found on gene expression for (D): F (1, 1792) = 749.5, p<0.0001, and a significant negative astrocyte effect was found on gene expression for (E): F (1, 2220) = 2427, p<0.0001. 2-way ANOVA performed on normalised gene expression levels (n=3 per condition). **F,G)** H3K27-Me3 ChIP peaks significantly increased (F) or decreased (G) in neurons by the presence of astrocytes whose location was in the gene or within 3 kb of the transcription start site were taken and the Log2 fold-change of mRNA expression plotted. A significant negative co-culture effect was found on gene expression for (F): F(1, 620) = 1280, p<0.0001, and a significant positive astrocyte effect was found on gene expression for (G): F(1, 708) = 144.1, p<0.0001. 2-way ANOVA performed on normalised gene expression levels (n=3 per condition). **H)** RNA-seq was performed on astrocyte-free cortical neuronal cultures (N=3) at DIV8 and DIV15. Genes with a DEseq2 fold-change of ≥1.5 fold up (red) or down (blue) due to astrocytes are highlighted (Benjamini Hochberg false discovery rate <0.05, average FPKM ≥1, here and throughout). **I)** Log2 DESeq2 fold-change of mRNA expression changes in developing cortical neurons due to age (DIV15 vs DIV8) was plotted against the effect of astrocyte co-culture. A weak negative correlation was observed (r= –0.102).

We first investigated the impact of astrocytes on the neuronal transcriptome. We performed RNA-seq on rat neuronal mono-culture and neuronal-astrocyte co-cultures (mixed species), and reads were allocated according to species using our Sargasso workflow ^8,15^, enabling the neuronal transcriptome to be specifically analysed in the co-culture as well as the mono-culture. We confirmed that the use of the Sargasso workflow has minimal effect on reported FPKM values compared to conventional (STAR) mapping (Fig. S1f, Pearson r correlation coefficient: 0.99). We found that cortical astrocytes induced 874 protein-coding genes and repressed 679 genes in cortical neurons (expression cut-off 1FPKM, fold-change ≥1.5X, Benjamini-Hochberg corrected *p*-value <0.05), indicative of a profound influence of astrocytes on developing neurons (Fig. 1b).

We also wanted to determine whether these astrocyte-dependent changes in gene expression are associated with an alteration of the neuronal epigenetic landscape. We chose two histone marks to study: Histone H3 acetylated on Lysine-9 (H3K27Ac) and Histone H3 tri-methylated on Lysine-9 (H3K27Me3) are known to be dynamically regulated and are associated with active open chromatin (H3K27Ac) and repressed/condensed chromatin (H3K27Me3) respectively. We performed ChIP-seq using antibodies against H3K27Ac and H3K27Me3 on both rat neuronal mono-cultures and mouse astrocyte/rat neuron co-cultures. ChIP-seq reads were allocated to species using our Sargasso workflow, enabling the profile of H3K27Ac and H3K27Me3 marks in neurons in the presence or absence of astrocytes to be defined without the imperfections or artefacts associated with physical sorting. The analysis revealed that astrocytes caused extensive changes to Histone H3 acetylation and methylation. In both cases over 3000 loci were significantly changed in neurons by the presence of astrocytes (EdgeR adjusted p-value <0.05, Fig. 1c).

We hypothesized that there would be a link between astrocyte-dependent H3K27 modification, and mRNA expression, so focussed on changes to H3K27 modification found in gene promoter regions up to 3kb from the transcription start site. Consistent with a link between H3K27 modification and gene expression, genes exhibiting an astrocyte-induced change in promoter H3K27 acetylation (up or down) showed a significant effect of astrocytes in mRNA levels in the same direction (Fig. 1d,e). Moreover, genes exhibiting an astrocyte-induced change in promoter H3K27 trimethylation showed a significant effect of astrocytes on mRNA levels in the opposite direction (Fig. 1f,g).

### Astrocytes cause the neuronal transcriptome to deviate from its intrinsic trajectory

We next wanted to determine whether the widespread astrocyte-induced changes in neuronal gene expression reflected a simple acceleration of the intrinsic developmental trajectory of the neuron transcriptome. If this were the case, then we would expect a correlation between the effect of astrocytes vs. the effect of culture time on neuronal gene expression. We performed RNA-seq on cultures of E20.5 rat cortical neurons that were maintained for 15 days, in the absence of astrocytes and under identical conditions as the DIV8 neurons. The purity of the neuronal population ((Fig. S1a-e) was further confirmed from our RNA-seq data: expression of rat *Gfap* and *Aldh1l1* in our neuron monocultures was 1000- and 400-fold fold lower respectively than in our mouse astrocyte monocultures.

Analysis of differential gene expression revealed that as expected, the transcriptome of DIV15 neurons showed widespread changes compared to DIV8, with 807 genes induced, and 448 genes repressed (Fig. 1h). We employed the same Sargasso workflow for consistency with the mixed species cultures, and confirmed that its use has minimal effect on reported fold changes compared to conventional (STAR) mapping (Fig. S1g, Pearson r correlation coefficient: 0.99)

We then plotted the Log2-fold change of all genes with culture time (DIV15 vs. DIV8) vs. Log2-fold change due to co-culture (with astrocytes vs. without). Strikingly, we observed an absence of correlation between intrinsic temporal changes in neuronal gene expression and astrocyte-induced changes in neuronal gene expression (Fig. 1i). This suggests that astrocytes induce specialised changes in developing cortical neurons that are not aligned with neurons’ intrinsic temporal gene expression trajectory, arguing against the concept that astrocytes only accelerate intrinsic neuronal maturation.

### Ontological analysis of astrocyte-dependent neuronal genes

To gain insight into potential functional consequences of astrocytes on developing cortical neurons, we performed ontological analysis on genes controlled in neurons by astrocytes (DIV8 co-culture vs. mono-culture). Genes induced by astrocytes in neurons were enriched (>4-fold, p<0.05) in GO Biological Processes associated with neurite outgrowth, axon guidance, and associated signal transduction pathways such including Semaphorin-, Kit- and Notch-signaling (Fig. 2a). Genes associated with cholesterol biosynthesis were also enriched, noteworthy since cholesterol plays an important role in dendrite formation and axon guidance. GO Molecular Function terms centred on a variety of signal transduction pathways, as well as voltage-dependent ion channels, many involved in action potential firing (Fig. 2b). Thus, astrocytes induce genes in neurons whose function would be expected to promote neurite outgrowth and influence the expression profile of receptors and ligands with the potential to determine future responses to extrinsic signals. We also investigated whether astrocyte-induced neuronal genes showed any enrichment for disease risk, since it may point to an importance for astrocyte-to-neuron signaling in the aetiology or progression of various brain disorders. Using published gene sets for disease risk ^16–21^, we found enrichment in the set of astrocyte-induced neuronal genes for genes associated with schizophrenia, epilepsy, attention deficit hyperactivity disorder (ADHD) and Alzheimer’s disease (AD) risk, but not autism spectrum disorder (ASD) or major depressive disorder (MDD, Fig. 2i).

**Figure 2.**
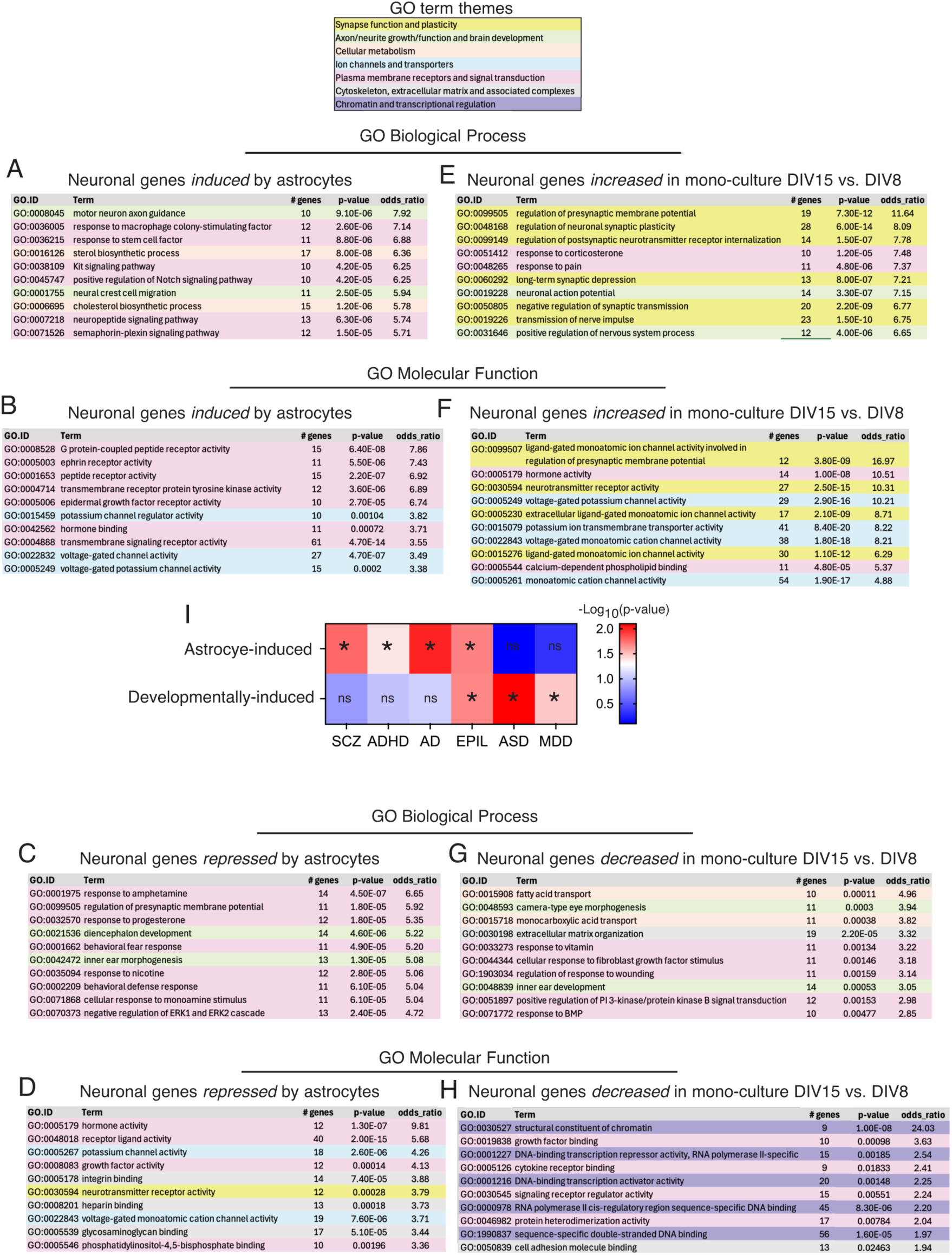
Ontological analysis of genes regulated as indicated by astrocytes (left) or by culture time period (right). **A-H)** GO terms are colour-coded by thematic area as indicated in the GO term themes box at the top of the figure. I) Genes up-regulated ≥1.5 fold by astrocytes (upper) or by developmental stage in mono-culture (lower) were tested for enrichment for genes associated with risk for the indicated disorders (see main text) *p=0.0169, 0.042, 0.0102, 0.0204 (upper), 0.0211, 0.0004, 0.0338 (lower), Fisher’s exact test.

Among genes in neurons whose expression is down-regulated by astrocytes, terms are also dominated by signal transduction Biological Process and Molecular Function gene sets, again suggestive of altered status of signaling pathways induced by astrocytes. Potassium channel terms were also prominent among down-regulated genes, including voltage-dependent ones as well as non-voltage activated channels such as those encoding *Kcnj3* (Kir3.1), *Kcnj4* (Kir2.3), *Kcnj2* (Kir2.1) and *Kcnj9* (Kir3.3), which can control neuronal intrinsic properties (Fig. 2c,d).

We also performed a GO analysis on the genes that are regulated as part of neurons’ intrinsic developmental trajectory (mono-cultures: DIV15 vs. DIV8). Genes up-regulated during development in an astrocyte-independent manner were enriched in multiple GO Biological Processes associated with synaptic transmission and synaptic plasticity (Fig. 2e). Moreover, Molecular Function terms centred on ion channel function, and while voltage-dependent ion channel terms were prominent (similar to the effect of astrocytes), so were neurotransmitter receptor gated channel terms such as (Fig. 2f). Given that the gene sets regulated by co-culture and regulated by development are very different (Fig. 1i), it is not surprising that they are enriched in different GO terms, as well as with regard to enrichment in disease risk genes, where enrichment in ASD and MDD risk was observed (unlike astrocyte-induced genes) and not significant for schizophrenia, ADHD, or AD risk (also unlike astrocyte-induced genes). Only epilepsy risk genes were enriched in both gene sets (Fig. 1i). Collectively this analysis underlines the differences between astrocyte-induced changes and neurons’ own intrinsic developmental trajectory.

### Astrocytes can induce network activity via the control of intrinsic properties

We next examined spontaneous activity of DIV8 cortical neurons maintained in the presence or absence of astrocytes, well established to be boosted by astrocytes in RGCs ^22^. We first performed Ca^2+^ imaging of neuronal mono- and co-cultures. Neurons co-cultured with astrocytes exhibited large and frequent spontaneous Ca^2+^ transients (Fig. 3a, b). The transients were synchronous across the field of view and were abolished by TTX treatment, consistent with them being due to action potential firing activity synchronously across the network. In contrast, astrocyte-free neuronal mono-cultures exhibited weak or non-existent spontaneous Ca^2+^ transients (Fig. 3a,b). We further investigated neuronal spontaneous activity in mono- and co-cultures by measuring spontaneous EPSCs in whole cell voltage-clamp. Consistent with the Ca^2+^ imaging data, neurons showed evidence of much larger synaptic activity in the presence of astrocytes. While the number of spontaneous events recorded were similar (Fig. 3c), the net excitatory current was nearly ten times higher in the presence of astrocytes than in mono-cultures (Fig. 3d). Thus, cortical astrocytes profoundly influence the electrophysiological output of developing cortical neuronal networks.

**Figure. 3.**
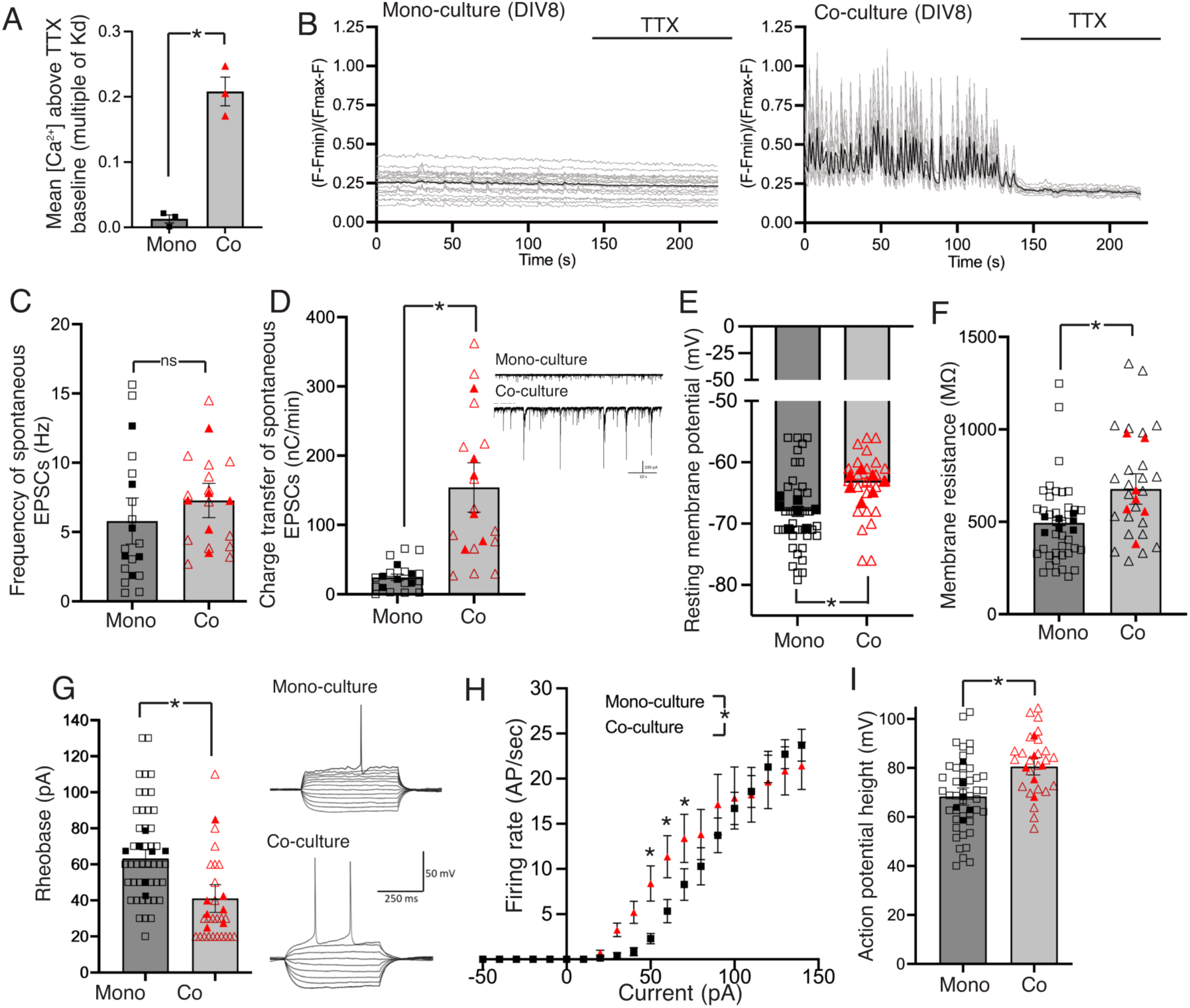
Astrocytes control the intrinsic excitability of developing cortical neurons. **A)** Fluo-3 Ca^2+^ imaging was performed on mono- and co-cultures at DIV8, with spontaneous transients recorded for a minimum of 100s before TTX (1 µM) added to block firing activity. [Ca^2+^] as a multiple of the indicators Kd (F-Fmin)/(Fmax-F) was calculated and the mean value over 100 s calculated, from which the mean [Ca^2+^] over the last 30s of imaging in the presence of TTX. 10-20 cells per coverslip were analysed and the mean taken as a single data point. t=8.524, df=4, p=0.001 (unpaired t-test, n=4). **B)** Example traces from a single experiment showing individual cells (light grey) and the mean (black). **C, D)** In DIV8 mono- and co-cultured cortical neurons spontaneous EPSCs were recorded and their frequency (C) and total charge transfer measured, with example traces in (D). C: t=0.7061, df=10, p=0.496; D: t=3.591, df=10, p=0.005 (nested t-test, n=6). **E-G)** Intrinsic properties of DIV8 cortical neurons measured. E (resting membrane potential): t=2.702, df=12, p=0.019. F (membrane resistance): t=2.576, df=12, p=0.024. G (rheobase): t=4.239, df=12, p=0.001. Nested- t-test (n=7). Inset to (J) shows example trace when calculating the rheobase. **H)** Relationship between current injection and action potential frequency (F/I curve). We applied 10 pA steps of 500 ms duration, at 5 s intervals. Astrocyte effect: F (1, 239) = 6.595, p=0.011 (2-way ANOVA), p=001, 0.014, 0.030 (Fishers post-hoc test). N=7. **I)** Effect of astrocytes on action potential height. t=2.409, df=10, p=0.037 (nested t-test, n=6).

To determine the synaptic contribution to this enhanced firing activity we measured excitatory synaptic function by analysing miniature EPSCs (mEPSCs) at both DIV8 and DIV15, in the presence and absence of astrocytes. We observed that mEPSC frequency increased from DIV8 to DIV15 (Fig. S2a, c). However, mEPSC frequency and size was unaffected by co-culture with astrocytes at either developmental stage assessed by both a 2-way ANOVA and a hierarchical nested t-test, as well as a paired t-test to allow for week-to-week variation in mEPSC culture frequency (Fig. S2a-f). Since there is precedent for cortical neuronal synaptogenesis being independent of astrocyte-released factors identified using the RGCs ^12,23^, we studied the influence of cortical astrocytes on RGCs. We cultured RGCs under the same conditions as cortical neurons, finding that cortical astrocytes strongly increased RGC mEPSC frequency (Fig. S2g, i, mEPSC size was unaltered, Fig. S2h) consistent with prior studies ^11–14,22^. We confirmed the absence of an effect of astrocytes on cortical neuronal synaptogenesis by immunohistochemical analysis of synapse density (pre/post synaptic marker colocalization) and spine density across defined dendritic sections (Fig. S2j, k). We also measured whole-cell AMPA and NMDA receptor currents and found them to be unchanged by the presence of astrocytes (Fig. S2l, m). Additionally, we assessed the influence of astrocytes on the developmental maturation of AMPA receptor composition, characterised by a reduction in GluA2-lacking AMPA receptors, and measured by the sensitivity of currents to (NASPM, 1-Naphthyl acetyl spermine trihydrochloride). We observed a developmental reduction in the NASPM sensitivity of AMPAR currents but not an influence of astrocyte presence (Fig. S2n, o). Thus, in our experimental system, both excitatory synaptogenesis and maturation of AMPA receptors in cortical neurons appears to proceed normally in the absence of astrocytes. Moreover, while blocking inhibitory neurotransmission with the GABA_A_ receptor antagonist gabazine increased sEPSC frequency (Fig. S2p), we still observed an astrocyte-dependent increase in excitatory charge transfer in the presence of gabazine (Fig. S2q), pointing to a non-synaptic mechanism of elevated network activity.

A key determinant of neuronal firing activity is the intrinsic excitability of the neuron, and so we measured a variety of standard parameters used to assess this. We observed that at DIV8, the presence of astrocytes caused cortical neurons to have a more depolarized resting membrane potential (Fig. 3e) and higher membrane resistance (Fig. 3f). A depolarized resting membrane potential will mean that a smaller voltage change will be required to initiate an action potential, and a higher membrane resistance means that a smaller current will be needed to achieve the necessary voltage change. Consistent with this we observed a lower rheobase in the presence of astrocytes (Fig. 3g), which is the minimum amount of current needed to trigger an action potential. Moreover, the relationship between current injection and action potential frequency (the F-I curve) of neurons cultured with astrocytes was strongly shifted to the left, with any given current eliciting a higher rate of AP firing that in mono-cultured neurons (Fig. 3h), and action potential height was slighter higher in the co-cultures (Fig. 3i). Thus, cortical astrocytes affect cortical neuronal intrinsic properties with the effect of increasing their excitability, representing a novel non-synaptic mechanism by which astrocytes can influence the output of developing cortical neurons.

### Astrocytes control the intrinsic excitability of developing cortical neurons via Kir suppression

We next investigated the basis for the astrocyte-dependent changes in neuronal excitability. We examined our transcriptomic data to identify gene expression changes that had the potential to alter neuronal excitability. Inwardly rectifying K^+^ channels (Kirs) are known to play a key role in determining the excitability of neurons ^24^, so it was notable that the four highest expressed genes encoding inwardly rectifying K^+^ channels, *Kcnj3* (Kir3.1), *Kcnj4* (Kir2.3), *Kcnj2* (Kir2.1) and *Kcnj9* (Kir3.3) were all significantly repressed in neurons by astrocytes (by 73%, 62%, 40%, and 36% respectively, Fig. 4a).

**Figure 4.**
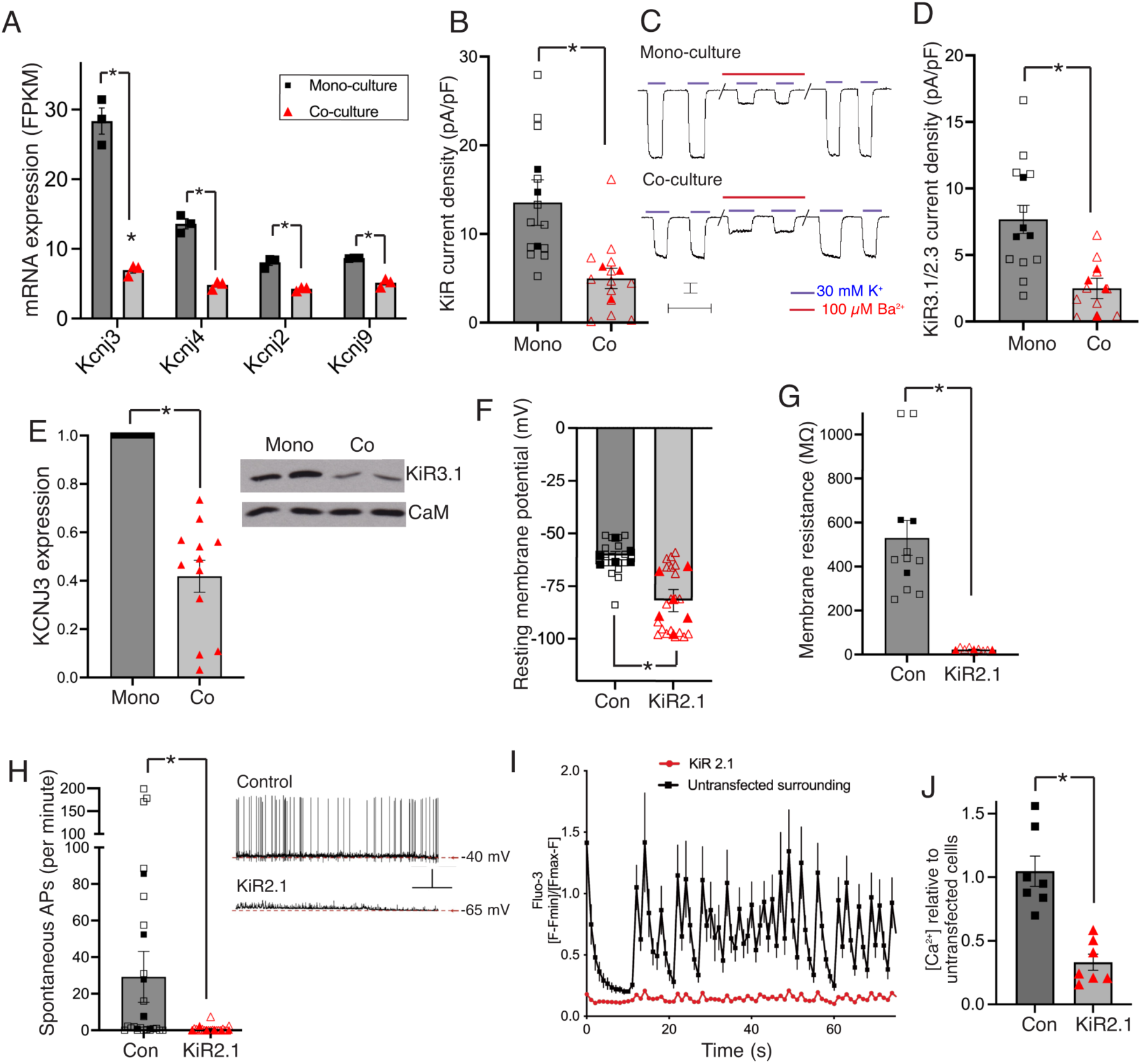
Astrocytes down-regulate neuronal expression of inwardly-rectifying potassium channels, determinants of intrinsic excitability. **A)** Influence of astrocytes on neuronal expression of the indicated *Kcnj* genes, taken from our RNA-seq data. *p= 6.46E-168, 5.54E-52, 2.66E-24, 4.67E-09 (Benjamini Hochberg adjusted p-value within DESeq2). **B)** Neuronal potassium channel currents (co-vs mono-culture) recorded before and after application of the pan-Kir antagonist BaCl_2_ to define whole-cell Kir current density: t=3.080, df=4, p=0.037 (nested t-test, n=3). **C)** Example traces showing K+-induced currents before, during and after BaCl_2_ treatment. Scale bars: 100 pA, 5 s. **D)** Potassium channel currents recorded before and after application of antagonists of the two highest expressed Kir genes Kcnj3/Kir3.1 (150 nM tertiapin Q) and Kcnj4/Kir2.3 (20 µM ML 133). Methodology as per (B): t=3.262, df=6, p=0.017 (nested t-test, n=4). **E)** Western blot analysis of Kir3.1 expression in the indicated cultures at DIV8. t=8.768, df=22, p<0.0001 (n=1). **F)** Resting membrane potential of Kir2.1 over-expressing neurons (co-cultured with astrocytes), compared to control (ß-globin-transfected): t=3.640, df=10, p=0.005 (nested t-test, n=6). An mCherry plasmid was co-transfection marker was employed. **G)** Membrane resistance measured in co-cultured Kir2.1 vs. control-transfected neurons: t=4.869, df=4, p=0.008 (nested t-test, n=3). **H)** Spontaneous firing rate in Kir2.1 vs. control-transfected neurons (in astrocyte co-culture): t=2.362, df=10, p=0.040 (nested t-test, n=6). Scale bars: 20 mV, 20 ms. **I, J)** Fluo-3 Ca^2+^ imaging was performed on co-cultures at DIV8 transfected with a plasmid encoding either Kir2.1 or ß-globin, plus a mCherry co-transfection marker. Spontaneous transients were recorded for a minimum of 50s [Ca^2+^] as a multiple of the indicator’s Kd (F-Fmin)/(Fmax-F) and the mean value taken relative to the level in surrounding un-transfected neurons. t=5.319, df=12, p=0.0002 (n=7, where more than one transfected cell was observed in a field of view the average of the cells was taken to give a single data point).

We then measured Kir channel activity directly by defining the change in current upon addition of a saturating concentration of the pan-Kir blocker Ba^2+^ (100 µM), and observed a reduction of 60 % in Kir current due to the presence of astrocytes (Fig. 4b,c). We also performed an identical experiment, replacing Ba^2+^ with inhibitors of Kir3.1 (tertiapin-Q) and Kir2.3 (ML133) (the two highest-expressed *Kcnj* genes) and observed a 70% reduction in combined Kir3.1/2.3 activity (Fig. 4d). We also confirmed astrocyte-induced repression of the highest expressing gene, *Kcnj3*/Kir3.1, by western blot (Fig. 4e). Thus, astrocytes reduce neuronal Kir activity in developing neurons through the coordinated repression of their genes. To determine the biological consequences of Kir hyper- or hypo-function in our system we first transfected neurons with a plasmid encoding one Kir (Kir2.1/*Kcnj2*) in co-cultures, with a mCherry transfection marker. We compared the impact of Kir2.1 expression to that of neurons transfected with a control plasmid encoding ß-globin. We found that Kir2.1 expression in co-cultured led to a hyperpolarization of their resting membrane potential (Fig. 4f), and decrease in membrane resistance (Fig. 4g). As expected, spontaneous firing activity was suppressed by Kir over-expression (Fig. 4h), as was activity-dependent spontaneous Ca^2+^ transients (Fig. 4i,j). Thus, artificially boosting Kir function in co-cultured neurons is sufficient to lower their intrinsic excitability and prevent spontaneous activity.

To determine whether a partial reduction in Kir currents (as seen in co-cultures) is sufficient to alter neuronal intrinsic properties in a manner similar to astrocyte co-culture, we took astrocyte-free cortical neuronal mono-cultures and measured intrinsic properties before and after application of inhibitors of Kir3.1 (tertiapin-Q) and Kir2.3 (ML133) each at a concentration approximating to their reported IC-50 concentrations. Application of these drugs significantly increased membrane resistance (Fig. 5a) reduced the rheobase (Fig. 5b) and induced a leftward shift in the F-I curve (Fig. 5c,d). Moreover, we observed the same results when employing the pan-Kir blocker BaCl_2_ at a concentration near its IC-50 (5 µM, Fig. 5e-g). We also observed an increase in net current influx into mono-cultured neurons due to spontaneous activity when 5 µM BaCl_2_ was applied (Fig. 5h). Thus, partial pharmacological blockade of Kirs in astrocyte-free neuronal cultures phenocopies the effect of astrocytes with regard to their effect on neuronal intrinsic properties, consistent with Kir repression as the mechanism behind the effects of cortical astrocytes on cortical neuronal network activity.

**Figure 5.**
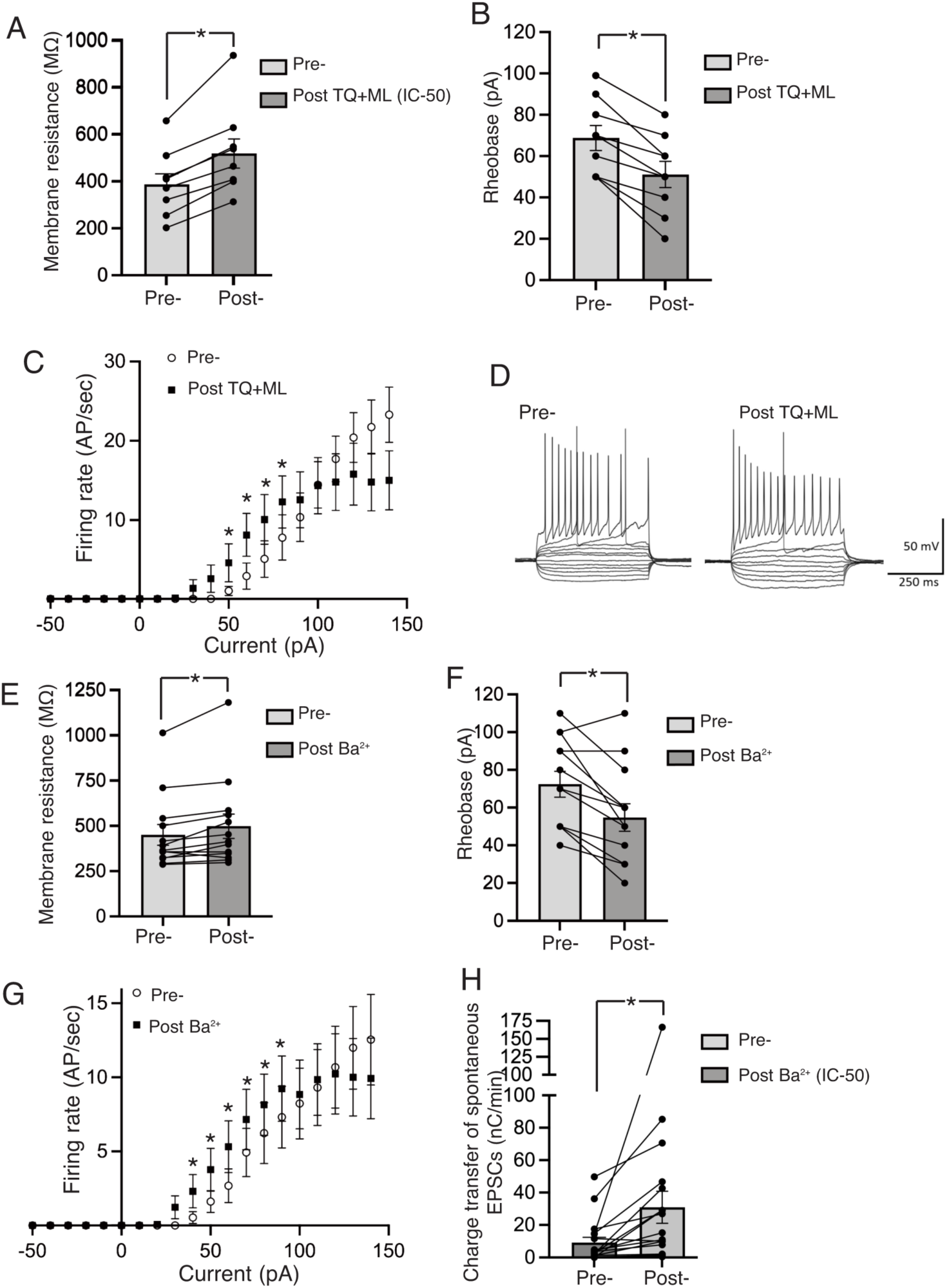
Partial inhibition of neuronal Kirs in neuronal mono-cultures mimics the effect of astrocytes on neuronal intrinsic properties. **A-D)** The intrinsic properties of neuronal mono-cultures were measured before and after the application of tertiapin Q (TQ, 15 nM) and ML133 (4 μM), inhibitors of Kir3.1 and Kir2.3/2.1 respectively. Non-saturating concentrating levels of these compounds were used to better mimic the suppressing effect of astrocytes. An increase in membrane resistance was observed (A), as was a decrease in the rheobase (B) and a left-ward shift of the F-I curve (C,D). A: t=6.604, df=8, p=0.0002 (two-tailed paired t-test, n=9); B: t=6.380, df=8, p=0.0002 (two-tailed paired t-test, n=9); C: F (19, 160) = 5.191, p-value<0.0001 (drug x current interaction, 2-way ANOVA), *p=0.026, 0.001, 0.002, 0.005 (Ficher’s post-hoc test, n=9). **E-G)** Experiments performed as in A-C except BaCl_2_ was applied (5 µM). E: t=3.197, df=12, p=0.008 (two-tailed paired t-test, n=13); F: t=4.153, df=12, p=0.0013 (two-tailed paired t-test, n=13). G: F (19, 240) = 3.169 P<0.0001 (drug x current interaction, 2-way ANOVA). *p=0.023, 0.037 (Fisher’s post-hoc test, n=9) **H)** Rate of spontaneous charge transfer upon application of 5 µM BaCl_2:_ t=2.567, df=17, p=0.020 (two-tailed paired t-test, n=9).

### Astrocytes release proteins to promote neuronal excitability

We wanted to know whether the effect of cortical astrocytes on Kir expression and on the neuronal transcriptome in general is mediated by contact-dependent signaling or via secreted molecules. We treated astrocyte-free rat cortical neuronal mono-cultures with astrocyte-conditioned medium (ACM) for 72h (from DIV0 to DIV3) and performed RNA-seq which revealed substantial changes in gene expression (Fig. 6a). In parallel we performed RNA-seq on rat neurons cultured with mouse astrocytes (also at DIV3) which also induced changes in gene expression when compared to neuronal mono-cultures (Fig. 6b). We then compared ACM-induced changes in gene expression with physical astrocyte co-culture-induced changes (Fig. 6c). We took genes significantly altered by either ACM treatment, or co-culture, or both and plotted ACM-induced changes against astrocyte co-culture-induced changes. We observed a good correlation (Fig. 6c), suggesting that the major influence of cortical astrocytes on cortical neuronal gene expression is exerted via secreted factors. Examination of *Kcnj* expression revealed that even after only 3 days, *Kcnj* expression is lowered both by astrocyte co-culture and ACM (Fig. 6d). This suggested to us that it should be possible to recapitulate the effect of astrocyte co-culture on neuronal intrinsic properties by culturing neuronal mono-cultures in ACM. Indeed, treatment of neuronal mono-cultures with ACM until DIV8 had a very similar effect to astrocyte co-culture: neuronal resting membrane potential was shifted to a more depolarized position (Fig. 6e), membrane resistance increased (Fig. 6f), rheobase decreased (Fig. 6g) and the F-I curve shifted leftwards (Fig. 6h)

**Figure 6.**
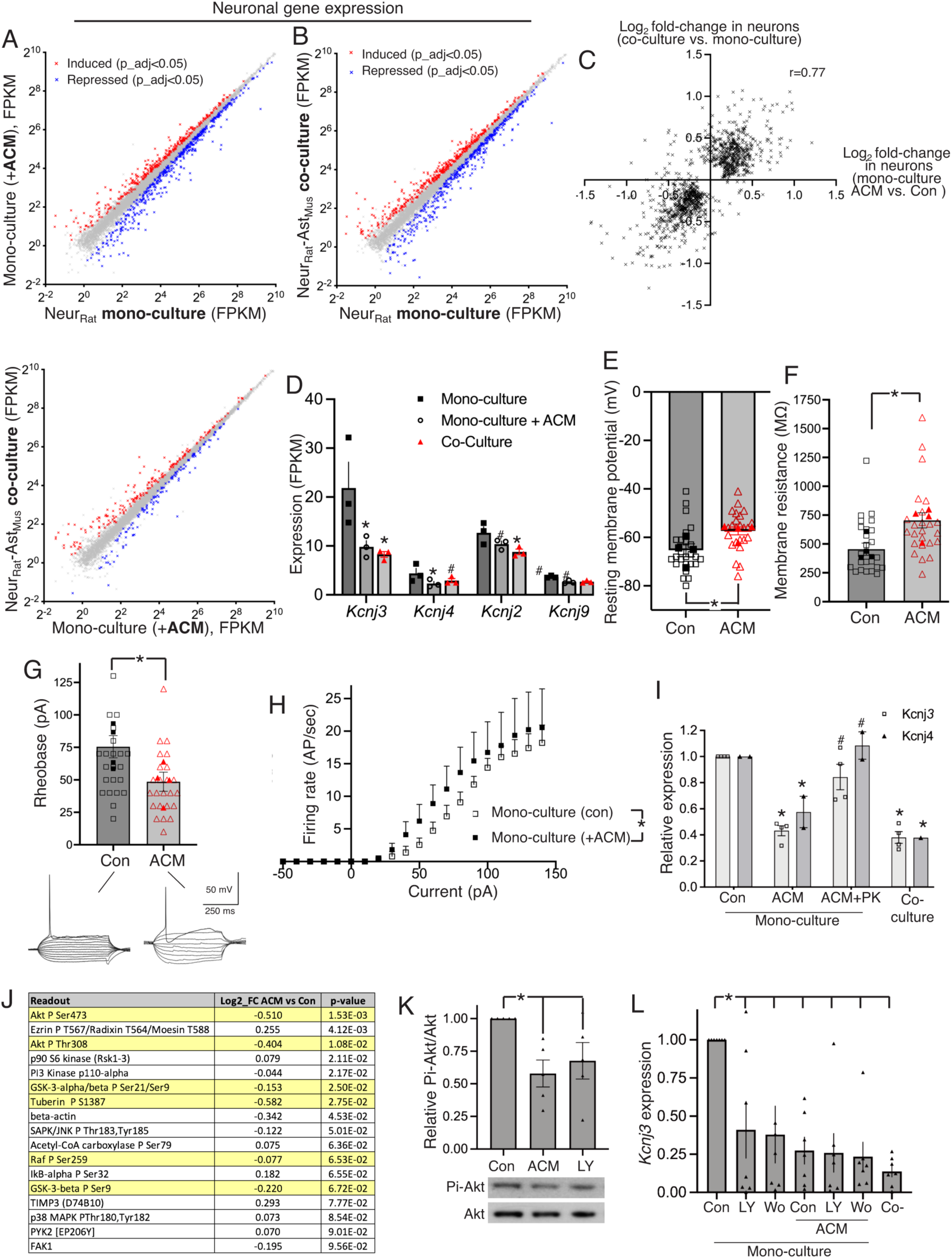
Astrocyte secreted proteins are sufficient to promote excitability of developing cortical neurons. **A)** RNA-seq was performed on neuronal mono-cultures treated for 72h (DIV0 to DIV3) with astrocyte conditioned medium (ACM) or control unconditioned medium. Genes induced (red) or repressed (blue) due to ACM treatment are highlighted (DEseq2 Benjamini Hochberg false discovery rate <0.05, average FPKM ≥1, n=3). **B)** RNA-seq was performed on the indicated cultures and both subjected to the same Sargasso workflow to assign reads to rat (neurons). Genes induced (red) or repressed (blue) due to ACM treatment are highlighted (DEseq2 Benjamini Hochberg false discovery rate <0.05, average FPKM ≥1, n=3). **C)** For all genes significantly changed by 72h ACM (A) or 72h co-culture in neurons (or both), the Log2 fold-change is compared and correlation coefficient (r=0.77) calculated. **D)** Expression of the indicated *Kcnj* genes taken from the RNA-seq data in (A) and (B). * DEseq2 Benjamini Hochberg adjusted p-value <0.05; #unadjusted p-value <0.05. **E-G)** effect of ACM treatment on neuronal mono-cultures. E (resting membrane potential): t=2.572, df=6 p=0.042; F (membrane resistance): t=3.250, df=6, p=0.018; G (rheobase): t=2.558, df=6, p=0.043 (nested t-tests, n=4). **H)** Effect of ACM on DIV8 mono-culture F-I curve. F (1, 120) = 5.272, p=0.023 (main effect of ACM, two-way ANOVA). **I)** ACM was treated ± protease K for 2 hrs prior to addition to neuronal mono-cultures and expression of Kcnj3 and Kcnj4 assessed. F (4, 19) = 23.01, p<0.0001 (2-way ANOVA). Sidak’s post-hoc test: *p values compared to control: 0.0002, 0.049, <0.0001, 0.016. # denotes p-values relative to ACM: 0.006, 0.016. **J)** Summary of results of a reverse phase protein array analysis of proteins and phospho-proteins modified by ACM treatment of mono-cultured neurons. Array components with a p-value of <0.1 are shown. Yellow highlights known targets of Akt. **K)** Western blot of phosphor- (Ser-473) Akt following ACM or LY294002 (5 µM) treatment of neurons. One way ANOVA (F(2,12)=4.47, p=0.029) followed by Fishers LSD: p=0.012, 0.042. **L)** Q-PCR analysis of Kcnj3 at DIV3 following the indicated treatments. All samples are mono-cultures except for the final co-culture sample (“Co-“). F = 5.52, p=0.0003 (One-way ANOVA) plus Dunnett’s post-hoc test: *p=0.007, 0.004, 0.0007, 0.0006, 0.0004, <0.0001, n=7. LY: 10 µM LY294002, Wo: 0.5 µM wortmannin.

We found that the influence of ACM on neuronal *Kcnj* expression was abolished by proteinase K treatment of the ACM, identifying the active factors as proteins (Fig. 6i). While efforts are ongoing to identify the factor(s) responsible, to begin to shed light on the mechanism behind astrocyte-dependent Kir repression we performed a reverse-phase protein array assay (RPPA) of extracts from neurons treated ± ACM to look for signalling pathways altered by ACM (the array contained many phospho-specific antibodies against signaling proteins). A prominent change was centred on the PI3K-Akt pathway: we observed that ACM treatment of mono-cultured neurons caused a reduction of Akt phosphorylation at both Ser-473 and Thr-308), markers of its activation (Fig. 6j). We also saw a reduction in levels of phospho-(Ser-9) GSK-3, phospho-(Ser-1387) Tuberin and phospho-(Ser-259) Raf, all known Akt targets.

We first confirmed a reduction in Akt phosphorylation (at serine-473 from the protein array data) upon ACM treatment of neurons by western blot (Fig. 6k), before investigating the effect of suppressing the Akt pathway on *Kcnj3* expression. Treatment of neuron mono-cultures with inhibitors of Akt’s upstream activator PI3K (wortmannin and LY294002) was sufficient to mimic the effect of ACM in repressing *Kcnj3* expression (Fig. 6m). Additionally, ACM treatment occluded the effects of wortmannin and LY294002 on Kcnj3 repression, suggestive of a common mechanism (Fig. 6l). Thus, astrocyte-mediated repression of Kir expression may be due to down-regulation of the PI3K-Akt pathway.

As a final investigation into neuronal *Kcnj3* regulation by astrocytes we wanted to confirm that the effect was not due to the neurons coming from a different species (rat) than the astrocytes (mouse). We performed single nucleus RNA-seq on rat and mouse cortical neurons cultured in the presence or absence of mouse astrocytes. While single-nucleus RNA-seq provides less depth of sequencing and reveals fewer differentially regulated genes, it nevertheless enables a direct comparison of the response of rat vs mouse neurons to mouse astrocytes. We identified genes induced or repressed >1.5-fold in rat neurons that have 1:1 mouse orthologs and are expressed in at least 50% of cells. We observed a good agreement between the genes induced or repressed in rat neurons, with the corresponding changes in mouse neurons (Fig. S3a-e), including the reduction of *Kcnj3*. Thus, the reduction of neuronal *Kcnj3*, and more broadly the changes observed in neurons, are not due to them being exposed to a different species of astrocyte.

### Astrocyte presence reveals neuronal homeostatic plasticity of intrinsic properties

Having established that astrocytes have profound effects on neuronal gene expression and intrinsic properties early in development, we wanted to know if this effect remains at later stages. We performed RNA-seq of co- vs mono-cultures at DIV15 and observed fewer gene expression changes than at DIV8 (261 genes altered >1.5 fold, Fig. 7a). Measurement of spontaneous neuronal activity revealed that the difference in spontaneous activity no longer reached significance (Fig. 7b), raising the possibility that long-term differences in activity-dependent signaling and gene expression may be leading to a homeostatic feedback control on neuronal activity and the properties of the neurons. To remove this potential effect, we treated both mono- and co-cultures with TTX for 48 h to abolish firing activity and once more compared co- vs mono-cultures at DIV15. In the presence of TTX, the effect of astrocytes was substantially greater than in its absence: 551 genes were significantly altered >1.5-fold (Fig. 7c). This suggested to us that neuronal activity may be influencing similar genes to astrocytes, but in the opposite direction. A comparison of DIV15 co-cultures ± TTX revealed a large number of genes subject to regulation by spontaneous activity in co-cultures (Fig. 7d). We then plotted the effect of astrocytes (Log2-fold change co-culture vs. mono-culture, both in the presence of TTX) against the effect of activity blockade (Log2-fold change co-culture ± TTX) and found a significant positive correlation between the effect of astrocytes and the effect of activity-blockade (Fig. 7e). Thus, at the genome-wide level, astrocytes appear to antagonize activity-dependent gene expression (despite positively influencing neuronal activity), potentially playing a role in bringing changes back to a baseline after activity has ceased, establishing a transcriptomic ‘ground state’.

**Figure 7.**
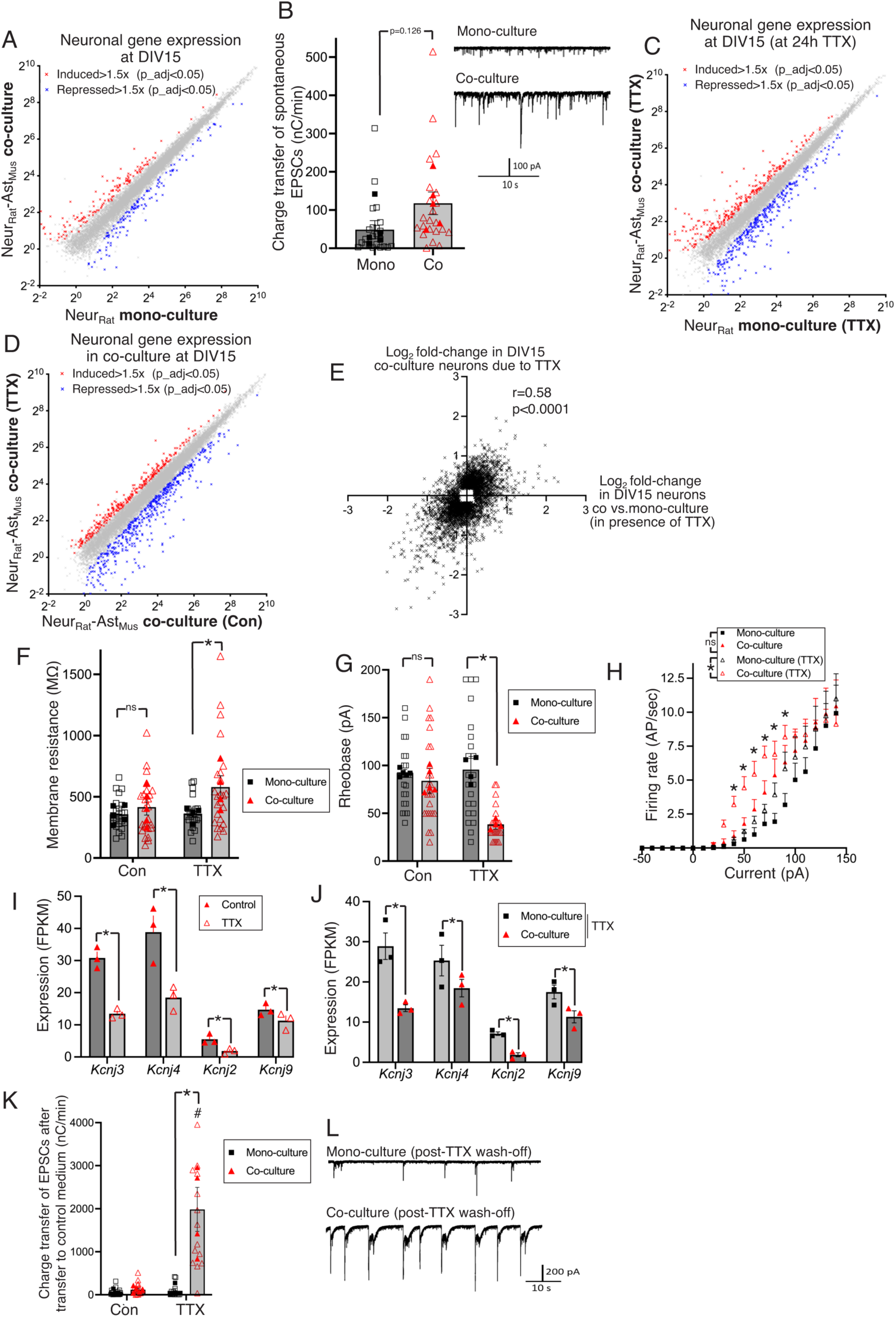
The pro-excitability effects of astrocytes on neurons cause an activation of neurons’ own homeostatic control of intrinsic properties. **A)** RNA-seq was performed on the indicated cultures and both subjected to the same Sargasso workflow to assign reads to rat (neurons). Genes with a DEseq2 fold-change of ≥1.5 fold up (red) or down (blue) due to astrocytes are highlighted (Benjamini Hochberg false discovery rate <0.05). **B)** Rate of spontaneous charge transfer due to neuronal activity in DIV15 mono and co-cultures. t=1.706, df=8, p=0.126, nested t-test, n=5. **C)** Experiment performed as in (A) except that both mono- and co-cultures were treated with TTX for 48h prior to harvesting. Genes with a DEseq2 fold-change of ≥1.5 fold up (265, red) or down (310, blue) due to astrocytes are highlighted (FDR <0.05). **D)** RNA-seq was performed on the indicated DIV15 co-cultures ± 48h TTX treatment prior to harvesting. Genes with a DEseq2 fold-change of ≥1.5 fold up (red) or down (blue) due to astrocytes are highlighted (FDR <0.05). **E)** A comparison of the influence of activity blockade (TTX) with the influence of astrocytes (co- vs mono-culture). The Log2 fold-change of genes significantly changed in either (or both) gene sets is shown and the correlation coefficient calculated. **F)** Influence of astrocytes on neuronal membrane resistance at DIV15 in the presence or absence of 48h TTX pre-treatment. F (1, 7) = 8.068, p=0.025 (astrocyte effect, 2-way ANOVA). *p=0.039, Sidak’s post-hoc test. **G)** Influence of astrocytes on neuronal rheobase at DIV15 in the presence or absence of 48h TTX pre-treatment. F (1, 7) = 47.11, p=0.0002 (astrocyte effect, two-way ANOVA). *p=0.0002 (Sidak’s post-hoc test). **H)** Influence of astrocytes on the neuronal F-I curve in the presence or absence of 48h TTX pre-treatment. F (1.516, 12.13) = 1.152, p=0.332 (ns, one-way ANOVA measuring interaction between the astrocyte effect and injected current, in the absence of TTX). F (19, 114) = 3.230, p<0.0001 (one-way ANOVA measuring interaction between the astrocyte effect and injected current, after homeostatic mechanisms had been re-set by 48h TTX pre-treatment). *p=0.015, 0.001, 0.0003, 0.006, 0.031 (Fishers post-hoc test, mono-culture vs. co-culture (after TTX pre-treatment). **I)** Effect of TTX on neuronal expression of the indicted *Kcnj* genes. Data taken from transcriptome-wide data in (D). *p=1.33E-05, 3.96E-05, 1.41E-08, 0.00361 (adjusted p-value created in DESeq2). **J)** Effect of astrocytes on expression of the indicated *Kcnj* genes after homeostatic mechanisms have been re-set by 24h TTX treatment. Data taken from transcriptome-wide data in (C). *p=7.99E-14, 1.67E-04, 1.97E-12, 2.81E-06 (Benjamini Hochberg adjusted p-value created in DESeq2). **K)** Rate of spontaneous charge transfer measured in both mono- and co-cultures after wash-off of TTX (compared to non TTX-treated pre-treated controls). F (1, 7) = 16.94, p=0.005 (astrocyte effect, 2-way ANOVA), *p=0.002 (Sidak’s post-hoc test). F (1, 7) = 19.68, p=0.003 (TTX pre-treatment effect, 2-way ANOVA), #p<0.0001 (Sidak’s post-hoc test). N=5 (control), 4 (TTX pre-incubation). **L)** Example traces from (K).

We also wanted to determine whether astrocytes still influence neuronal intrinsic properties and what role, if any, neuronal activity plays controlling these properties. Interestingly, we observed that at DIV15, co-cultured neurons have similar intrinsic properties to mono-cultured neurons: membrane resistance, rheobase and input-output curves were similar in co- and mono-cultures (Fig. 7f-h). This could mean that the effect of astrocytes is transient, however, another possibility is that activity-dependent homeostatic mechanisms are triggered to regulate neuronal excitability at DIV15. From at least DIV8 onwards, cortical neurons in the presence of astrocytes experience substantially more network activity that those maintained in the absence of astrocytes. This raises the possibility that the change in excitability in co-cultures between DIV8 and DIV15 is due to feedback inhibition of excitability.

To test this, we sought to reset any activity-dependent homeostatic mechanisms by blocking action potential firing for 48h using TTX treatment. Under these conditions, the differences in intrinsic properties between mono- and co-cultures was restored: a higher membrane resistance, lower rheobase, and a leftwards shift in the F-I curve (Fig. 7f-h). This demonstrates that astrocytes are still regulating neuronal intrinsic properties to promote excitability, but that their effects are countered by activity-dependent homeostatic feedback which lowers neuronal excitability. Examination of our RNA-seq data revealed that TTX treatment caused a reduction in expression of K*cnj3, Kcnj4, Kcn2* and *Kcnj9* in co-cultures (Fig. 7i). Moreover, in the presence of TTX, co-cultured neurons express lower *Kcnj3, Kcnj4, Kcnj2* and *Kcnj9* expression than mono-cultures at DIV15, as observed at DIV8 (Fig. 7j). Thus, when firing is blocked to reset homeostatic plasticity, *Kcnj* genes are suppressed and the effect of astrocytes on intrinsic properties becomes apparent. Consistent with the dramatic increase in excitability upon TTX pre-treatment of DIV15 co-cultures, placing TTX pre-treated neurons in control medium to observe spontaneous activity revealed a large difference in activity between mono- and co-cultures (Fig. 7k,l). In conclusion, our data supports a model whereby the influence of astrocytes on neuronal intrinsic properties pushes them into a zone of excitability where neurons’ activity-dependent homeostatic mechanisms regulate *Kcnj* expression which, in concert with other mechanisms ^25^, limit intrinsic excitability and prevent network hyperactivity.

## Discussion

For all stages of development except the very earliest, neurons are in close proximity to astrocytes. In maturity, neuronal function and survival relies on a combination of cell intrinsic mechanisms and non-cell-autonomous influences from glial cells, particularly astrocytes ^3,26–29^. During development, the influence of astrocytes on neurons has focussed on synaptogenesis. However, our analysis of astrocyte-to-neuron signaling here has revealed a strong influence of astrocytes on the epigenome and transcriptome, leading to the uncovering of an additional non-synaptic mechanism for influencing firing properties in developing cortical neurons.

### Astrocyte-dependent effects on the transcriptome of developing cortical neurons

Our previous study used the approach of mixed species RNA-seq ^15^ to define the widespread influence of cortical neurons on the cortical astrocyte transcriptome ^8^. Here we have studied the reciprocal influence of astrocytes on the neuronal transcriptome. Knowledge of the genes controlled provided us with hypotheses as to how astrocytes could regulate the intrinsic properties of neurons i.e. repression of Kir expression. Of course, among the hundreds of other genes controlled (both up, down and alternatively spliced) by astrocytes will be responsible for other functional changes induced in neurons by astrocytes, such as neurite outgrowth. Our study revealed a substantial number genes regulated by astrocytes in neurons whose function is associated with the processes of axon guidance, axogenesis and neuron differentiation, consistent with a role for astrocytes in neuronal maturation. Given the reciprocal nature of neurons and astrocytes on each other’s transcriptomes (this study and ^8^), it raises the question as to whether the influence of astrocytes on neurons is firstly dependent on the neurons themselves influencing astrocyte properties. While there is very likely to be some neuronal effects that do rely on neuron-to-astrocyte signaling, this is not universally the case. We found that conditioned medium from astrocyte mono-cultures (ACM) induces gene expression changes that correlate well with the influence of physical co-culture (Fig. 6c). Moreover, ACM was found to be sufficient to repress Kir gene expression and change the intrinsic properties of neurons (Fig. 6d-h). As such, in the case of astrocyte-dependent control of neuronal intrinsic properties both “naïve” mono-cultured astrocytes and co-cultured astrocytes have a similar effect. Thus, neuron-to-astrocyte signaling is not required for astrocytes to have the capacity to alter neuronal intrinsic properties.

Mechanistically, the primary responsible for mediating gene expression changes in response to astrocyte-derived signals await further investigation. The experiments using ACM point to a secreted factor, arguing against contact-dependent mechanisms. With the large number of genes up-and down-regulated it appears unlikely that a single factor is responsible, but factors capable of mimicking the effect of ACM at least in part are under investigation in the laboratory. This information is needed in order to define the importance of various astrocyte-induced gene expression changes in vivo, since astrocytes cannot be eliminated in the developing brain without deleterious consequences (unlike microglia ^30^). Instead, astrocyte-specific interference of the expression of putative factors would need to be achieved to define the in vivo impact.

### Astrocyte-dependent regulation of excitability in developing cortical neurons

We found that presence of astrocytes could dramatically increase activity across networks of cultured cortical neurons independently of synaptic changes. The excitatory synaptogenic potential of cortical astrocytes has been demonstrated many times using retinal ganglion cells (RGCs) as the target neurons ^9–14,23,31,32^. We were able to reproduce the strong effect of cortical astrocytes on RGC mEPSC frequency (Fig. 3k), but under our conditions we did not see a significant effect on intracortical neuronal functional excitatory synaptogenesis. It is therefore possible that neuronal type plays a role with regard to astrocyte-induced synaptogenesis. Interestingly, hevin is an astrocyte-secreted factor that promotes synaptogenesis in RGCs ^12^, but gain and loss of function studies showed that hevin signaling does not promote intracortical synapse number in vitro or in vivo ^23,33^. While most co-culture studies have employed RGCs, a study reported has increased FM-134 positive synaptic puncta in cortical neurons induced by astrocyte-conditioned medium ^34^, so the discrepancy may not lie in the neuronal type, but be due to differences in experimental conditions. While the reason for this would be interesting to determine, it is not central to the study which is a focus on non-synaptic effects.

The fact that neuronal activity could be induced independent of excitatory synaptic changes led to the uncovering of an additional mechanism whereby astrocytes could determine the electrophysiological output of developing neurons i.e. on controlling their intrinsic excitability. Astrocytes, or ACM, directly exert long-lasting control of cortical neurons’ own intrinsic excitability via altering neuronal gene expression. The coordinated down-regulation of neuronal Kir channels appears to be a major contributor to the alteration in neurons’ intrinsic properties. However, other gene expression changes may also contribute as other K^+^ channel genes are also altered at the transcriptional level, as are other ion channels. As with most cell types, astrocytes maintained in vitro have differences to those in vivo, although they have more in vivo-like morphology and properties when co-cultured with neurons as in this study ^5,6,8^. Nevertheless, future studies should address whether astrocytes exert a similar effect in vivo as in vitro. This may (for example) require the identification of the astrocyte-secreted factors that control neuronal intrinsic properties in vitro and generate astrocyte-specific knockouts of these factors to assess the influence on neuronal properties. More generally, the issue of why astrocytes are put in control of neuronal excitability in developing cortical neurons is a matter of conjecture. One possibility involves the role that astrocytes play at the synapse. Astrocytes are generated from neural precursor cells after neurons, but are essential for neurotransmitter uptake at the synapse, as well as spatial K+ buffering to clear the extracellular space of K^+^ during neuronal repolarization ^4,35,36^. Thus, astrocytes prevent the build-up of glutamate in and around excitatory synapses, and prevent neurons from chronically depolarizing by maintaining low extracellular K^+ 37^. As such, it might make teleological sense for neurons to remain relatively inactive until astrocytes are present to carry out the important tasks of neurotransmitter uptake and ion homeostasis. Elevated extracellular glutamate or chronic depolarization can be harmful to neurons, especially in allowing activation of extrasynaptic NMDA receptors and shut-off of activity-dependent neuroprotective pathways ^4,26,38–42^. More broadly, our study shows that the non-cell-autonomous influence of astrocytes on neuronal gene expression has a significant inverse correlation with the cell-autonomous effect of electrical activity on gene expression (Fig. 7e). Activity-dependent gene expression takes place in all mammalian neurons studied to date and is substantially conserved across evolution ^43,44^. It acts, via Ca^2+^ signaling, to promote metabolic, neuroprotective and developmental adaptions, as well as neuronal plasticity and homeostasis ^38,45–47^. The influence of astrocytes in tonically repressing activity-dependent gene expression programs (independent of their effects on neuronal activity-which in any case would exert effects in the opposite direction) requires further investigation, both in terms of molecular mechanism and biological reason, but may serve to promote the re-setting of activity-dependent genes back to baseline, improving the signal amplification of activity-induced dependent transcriptional changes.

### Astrocyte- and activity-dependent mechanisms combine to regulate neuronal excitability

It is well-established that neuronal activity (and by extension the networks of which they are part) are subject to activity-dependent homeostatic feedback control to maintain activity levels within acceptable limits, and involve control at both the synaptic level as well as intrinsic properties ^25,48–50^. Such homeostatic mechanisms can be reset by chronic blockade of firing activity, revealing the underlying basal state, and also reveal the extent to which homeostatic mechanisms have been invoked to modulate excitability. In this study we provide evidence that in the presence of astrocytes, spontaneous activity within the networks of cortical neurons is such that by DIV15 homeostatic mechanisms have been triggered which reduce neuronal excitability, since chronic activity blockade re-sets these changes and reveals an excitable ‘basal state’ which has a lower rheobase, lower membrane resistance, and left-shifted F-I curve. Moreover, TTX washout leads to strong rebound burst activity within the network, as a result of the high excitability of these ‘re-set’ neurons. This raises the possibility that different types of homeostatic plasticity (e.g. intrinsic plasticity vs. synaptic scaling ^51^) may have different importance in different cell types, or distinct thresholds for activation, a topic for future investigation.

In contrast to the situation in astrocyte-exposed cortical neurons, neurons maintained to DIV15 in the absence of astrocytes show no evidence of activity-dependent homeostatic regulation of neuronal excitability, since no effect of chronic activity blockade is observed on the rheobase, membrane resistance or F-I responses. Also, TTX washout does not result in rebound burst activity, since the intrinsic properties of the neurons have not changed. This can be explained by the fact that in the absence of astrocytes, DIV15 cortical neurons have low intrinsic excitability, not dissimilar (in terms of rheobase, membrane resistance and F-I responses) to astrocyte-exposed DIV15 neurons that are experiencing far higher levels of spontaneous activity, but which have activated excitability-repressing homeostatic mechanisms. As a result, activity levels in astrocyte-free cortical neurons are too low to invoke homeostatic pathways.

Collectively these observations are consistent with a model whereby cortical astrocytes provide constitutive, tonic, pro-excitatory signals to developing cortical neurons, at least in part by repressing Kir expression. The effect of this is to push developing cortical neurons into a zone of excitability that enables activity levels to rise to a point at which neurons’ own homeostatic mechanisms are triggered to keep network activity at appropriate levels. While activity-dependent homeostatic plasticity is multi-faceted ^25^, the dynamic activity-dependent induction of *Kcnj* gene expression suggests that it may contribute to this process, in direct opposition to the tonic astrocyte-dependent repression of *Kcnj* expression. In the scenario where neurons are maintained in the absence of astrocytes, cortical neurons’ weak intrinsic excitability is such that they do not reach this zone of excitability. Thus, astrocyte-to-neuron signaling may provide a floor below which intrinsic excitability cannot fall, ensuring that developing neurons remain capable of responding to inputs. It is an intriguing possibility that astrocytes play an important role in controlling neuronal excitability during the early critical periods of neuronal plasticity, potentially in the opening of this important developmental window ^52^.

A number of brain disorders have been associated with aberrant neural circuit activity, be at the level of firing rate, synchrony, oscillatory power or input response reliability ^53–55^. Such changes may be brought about by the effects of age, environment or gene mutations on the steady-state properties of the circuit, or indeed the capacity of these circuits to remain within normal parameters by a failure of homeostatic mechanisms. While astrocytes have long been known to play core roles in regulating the fidelity of synaptic transmission (such as through neurotransmitter uptake) and of ionic homeostasis of the extracellular milieu, their role in regulating neuronal gene expression that in turn influences circuit properties is not well understood. Our observation that risk genes for epilepsy, ADHD and schizophrenia are enriched in the gene sets controlled by astrocytes is interesting in this context as all are associated with aberrant neural circuit activity ^56–60^. In the context of Alzheimer’s disease, astrocyte-induced risk genes *Abca7* and *Mme* (neprilysin) are involved in Aß uptake and degradation ^61–63^. They form part of an enriched GO term among astrocyte-induced neuronal genes (‘amyloid-beta clearance by cellular catabolic process’), one of two enriched AD-related gene sets, the other being ‘cellular response to amyloid-beta’. It remains to be investigated whether astrocytes can functionally control ß-amyloidopathies through their effect on neuronal gene expression.

Overall, future work will be needed to fully address the implications of the astrocytic control of neuronal gene expression in vivo, in both health and disease. Such effects are likely to go beyond the electrophysiology of neural circuits, potentially extending to structural and metabolic changes, as well as susceptibility to disease-causing agents. Moreover, given the heterogeneity of both neurons and astrocytes, region and circuit-specific differences may emerge also.

## Acknowledgements

This work was funded by the UK Dementia Research Institute whose principal funder is the UK Medical Research Council, the Dunhill Trust, the Simons Initiative for the Developing Brain, and Wellcome Award 208402/Z/17.

## Author Contribution Statement

GEH conceived the project, and directed the project with DJAW, PCK, MD and OD. AT, JQ, KW, KH performed the experiments. OD, XH, KE and DV performed bioinformatic analysis. GEH wrote the manuscript. Other authors provided critical feedback on the manuscript.

## Materials and Methods

### Cortical tissue culture

All procedures described were performed in the University of Edinburgh in compliance with the UK Animals (Scientific Procedures) Act 1986 and University of Edinburgh regulations, approved by Edinburgh Local Ethical Review Board, conform to ARRIVE guidelines and carried out under project license numbers 70/9008, P1351480E and PP2262369. Mice were group-housed in environmentally-enriched cages within humidity and temperature controlled rooms, with a 12-hour light dark cycle with free access to food and water. Primary cell cultures of cortical neurons and cortical astrocytes were generated from tissue collected from embryonic day 17.5 CD1 mice and embryonic day 20.5 Sprague-Dawley rats. Briefly, embryos were euthanized by decapitation, their brains removed, and cortices dissected out into dissociation media (DM+K: 81.8 mM Na_2_SO_4_, 30 mM K_2_SO_4_, 5.84 mM MgCl_2_, 252 μM CaCl_2_, 1 mM HEPES, 0.1% Phenol Red, 20 mM glucose and 1 mM kyurenic acid). Cortices were placed in round bottom culture tubes (Corning) and digested in DM+K containing 20 units/mL of L-cysteine-activated papain enzyme (Worthington Biochemical Corporation) for 40 minutes at 37°C, shaking every 10 minutes, with fresh papain/DM+K solution added halfway through. Following digestion cortices were washed twice with DM+K, and twice with 1% NBA media, containing Neurobasal-A (Gibco), Anti-Anti (anti-bacterial/antimycotic, Invitrogen), B27 supplement (Life Technologies), glutamine (1 mM, Sigma) and 1% rat serum (Envigo). Cortices were then placed in a fresh 2 mL of 1% NBA and triturated gently using a 5 mL serological pipette on a low speed pipette gun (approximately 50 times). A further 2 mL of 1% NBA was then added to the tube, and after letting the tissue settle for a minute, the top 2 mL of suspension was collected and placed in a fresh tube. The remaining suspension was again triturated. If no tissue chunks remained it was added to the collected suspension, if not then a further 2 mL of 1% NBA was added, and the process repeated until all cells were completely dissociated. The dissociated suspension was then topped up to 10 mL with 1% NBA, and added to Opti-MEM (Gibco) solution supplemented with glucose (20 mM¬, Sigma) and Anti-Anti, at a concentration of 1 rat cortex/14 mL solution, and 1 mouse cortex/7 mL solution (giving an approximate concentration of 1,000,000 cells/mL). For neuronal cultures, 0.5 mL cell suspension (500,000 cells/well) was plated down onto glass coverslips (VWR) that had been placed into the wells of 24-well plates (Greiner), and pre-coated with poly-d-lysine (5 μg/mL, Sigma) and Laminin (13 μg/mL, Roche). For co-culture experiments, the coverslips had been pre-seeded with astrocytes 72 hours prior. The plates were then incubated at 37°C with 5% CO2 for two hours, before aspirating the Opti-MEM solution, and feeding with 1 mL 1% NBA containing 4.8 μM cytosine arabinoside (AraC). On day in vitro (DIV) 4 the wells were topped up with an additional 1 mL of fresh feeding media. For astrocyte cultures, 10 mL of cell suspension (∼130,000 cells per cm^2^) was pipetted into a 75 cm^2^ cell culture flask (Greiner) pre-coated with poly-d-lysine (5 μg/mL, Sigma). The plates were incubated at 37 °C with 5% CO_2_ for two hours, before aspirating the Opti-MEM solution, and feeding with 20 mL of DMEM solution (Dulbecco’s Modified Eagle Medium with high glucose, L-glutamine, sodium pyruvate and phenol-red, Gibco) that was supplemented with Anti-Anti and 10% fetal bovine serum (Gibco). On DIV4, 10 mL of DMEM was removed and a fresh 10 mL added. To generate pure astrocytic cultures the flasks were passaged on DIV7 and DIV11. Briefly, cells were washed with phosphate buffered solution (PBS, Gibco), and then incubated in 5 mL of 0.05% trypsin-EDTA (Gibco) at 37 °C for 5 minutes. The flasks were then gently rocked to ensure all cells were detached from the flask, collected into a 15 mL falcon tube with 5 mL of DMEM, and centrifuged at 800 RPM for 4 minutes. The supernatant was aspirated, and the cells were dissociated and collected into fresh DMEM at a concentration of 1x 75 cm^2^ flask of cells into 50 mL media. This suspension was then divided between three new poly-d-lysine coated 75 cm^2^ flasks (15 mL per plate). On DIV11 the process was repeated, but instead plating the astrocyte suspension onto coated coverslip-containing 24-well plates at a concentration of 1 mL/well (at a cell density of ∼100,000 astrocytes per coverslip). Nota bene, as astrocytes are proliferative cells, the final density of astrocytes per coverslip at the time of recording will be greater than the density at plate down. On (neuronal) DIV7 a 50% media exchange was done to feed the cells. For cells that were to be grown to DIV15, on DIV9, 11 and 13/14, 50% media exchanges were carried out using 0% NBA media supplemented with 10 mM glucose. Cells grown to DIV21+ followed the same feeding schedule, except on DIV13/14 a 50% media exchange was done with transfection media (TMITS) containing (in mM): 114 NaCl, 5.3 KCl, 1 MgCl_2_, 2 CaCl_2_, 10 HEPES, 1 glycine, 30 glucose, 0.5 sodium pyruvate, 0.2% NaHCO_3_, 0.001% phenol red, 10% Minimum Essential Media (-glutamine, Gibco), 1% Anti-Anti and 1% insulin-transferrin-selenium (100x, Invitrogen). Fifty percent media exchanges with TMITS were repeated on DIV17 and DIV21.

### Retinal ganglion cell culture

Preparation of retinal ganglion cells (RGC) was done following the Cold Spring Harbor Laboratory protocol ^64^, with minor adjustments. Postnatal day 5-6 rat pups were euthanized by decapitation before the eyeballs were enucleated into chilled DM+K media and kept on ice. Retinas were dissected from the eye in chilled DM+K, by first making an incision into the front of the eye, removing the lens and vitreous matter and then carefully tearing the sclera away from the retina. Retinas were kept in ice cold DM+K until all dissections were finished, before being transferred to a round bottomed culture tube containing DM+K with 20 units/mL of L-cysteine-activated papain enzyme and deoxyribonuclease I (DNAse; 0.07%, Worthington) and incubated for 30 minutes at 37 °C. The enzyme was removed and replaced with 2 mL of low ovomucoid solution (low-ovo), consisting of a 1.5% BSA (Sigma), 1.5% trypsin inhibitor (Worthington) and 0.07% DNAse D-PBS (Gibco) solution. A rabbit anti-rat macrophage antibody (80 μL, Cedarlane) was then added to the remaining 6 mL of low-ovo solution and mixed thoroughly. The low-ovo solution was aspirated from the retinas and a fresh 1 mL of the remaining low-ovo/anti-rat macrophage solution was added. The retinas were then gently triturated four times with a 1 mL pipette before adding a further 1 mL of low-ovo solution, with the top 1 mL then collected into a fresh falcon tube after the cells had settled. This was repeated until all of the 6 mL of low-ovo solution was used. The 6 mL of dissociated retinal cells was left to incubate at room temperature for 10 minutes to allow binding of the anti-rat macrophage antibody before being centrifuged for 12 min at 1,000 RPM. The supernatant was removed, and the cells were re-suspended in 6 mL of a high-ovomucoid D-PBS solution (high-ovo) containing 3% BSA and 3% trypsin inhibitor, before being returned to the centrifuge for a further 12 minutes. The high-ovo solution was removed and replaced with panning buffer made from D-PBS containing 0.01% BSA and insulin (5 μg/mL, Sigma). The panning buffer cell suspension was then passed through a 20-micron filter (pluriSelect). The filtered cells were poured onto a 15 cm negative panning dish pre-coated with goat anti-rabbit IgG (H + L) (Jackson Immunoresearch) and left at room temperature for 15 minutes, before transferring the solution of non-adherent cells onto a second pre-coated negative panning plate for a further 45 minutes, shaking every 15 minutes. The macrophage depleted retina cell solution was then transferred onto 10 cm positive panning plate, pre-coated with goat anti-mouse IgG (H + L) (Jackson Immunoresearch) and an anti-Thy1.1 antibody (clone OX-7, Sigma) and left for another 45 minutes, shaking every 15 minutes. The positive plate was aspirated and gently washed several times with D-PBS so only adherent cells remained. The plate was rinsed with calcium-free EBBS (Sigma) before incubating in 0.05% trypsin-EDTA for approximately 4 minutes at 37 °C until adherent cells shook off. The solution was collected into a falcon tube with D-PBS containing 30% fetal bovine serum (Gibco) and centrifuged at 1,000 RPM for 12 minutes. The supernatant was aspirated, and the retinal ganglion cells were resuspended in 1 mL 1% NBA and counted. They were then plated at a density of 50,000-100,000 cells per coverslip in RGC growth media consisting of 1% NBA with added forskolin (4.2 μL, Tocris), BDNF (50 ng/mL) and CNTF (10 ng/mL), onto coverslips in 24 well plates pre-coated with poly-d-lysine and laminin, or else onto coverslips pre-seeded with a bed of mouse astrocytes. Cells were fed with fresh RGC growth media every four days.

### Transfection

Neurons were transfected in TMITS media using Lipofectamine 2000 (2.33 μL/well, 1 μg/mL, Life Technologies), with a total plasmid concentration of 0.60 – 0.65 μg/mL. The Lipofectamine 2000/plasmid mix was incubated at room temperature for 20 minutes, before adding TMITS to give a total volume of 300 μL/well. Media from the wells to be transfected was removed (and saved aside if from neurons) before adding 300 μL of plasmid solution to each well. For astrocyte transfections, cells were transfected 48 hours after their plate down onto coverslips (before neuronal co-culture) and incubated with the plasmid mixture for 45 minutes at 37 °C, before being aspirated and fed with 1 mL DMEM media. For neuronal transfections, cells were co-transfected with green fluorescent protein (GFP) or mCherry on DIV4 and left in the plasmid mixture for 2 hours at 37 °C before aspiration. They were then fed with saved media and topped up with an additional 1 mL of fresh 1% NBA (+AraC) media. The plasmid encoding Kir2.1 was a gift from Matthew Nolan.

### Electrophysiological recordings

Standard electrophysiological recordings were performed in an external solution of artificial cerebrospinal fluid (aCSF) containing (in mM, all Sigma): 150 NaCl, 2.8 KCl, 10 Na-HEPES, 2 CaCl_2_, 1 MgCl_2_ & 10 glucose, and pH adjusted to 7.3 with NaOH. For RGC recordings, the aCSF contained (in mM): 150 NaCl, 2.8 KCl, 10 Na-HEPES, 2.5 CaCl_2_, 2 MgCl_2_ & 10 glucose, pH adjusted to 7.3 with NaOH. For neuronal current-clamp and intrinsic property recordings a K-gluconate internal solution was used consisting of (in mM): 130 K-gluconate, 4 glucose, 10 Na HEPES, 0.1 EGTA, 0.025 CaCl_2_, 20 sucrose, pH adjusted to 7.3 with KOH. For neuronal and RGC voltage-clamp recordings of spontaneous activity and mEPSCs, a Cs gluconate internal solution with 8 mM Cl^-^ was used, consisting of (in mM): 140 Cs gluconate, 3 CsCl, 0.2 EGTA, 10 HEPES, 5 QX-314 chloride, 2 Mg-ATP, 2 Na-ATP, 0.3 Na_2_GTP, 10 phosphocreatine, pH adjusted to ∼7.4 with CsOH. All recordings were performed at room temperature (20-23°C), using an Axopatch 200B amplifier (Molecular Devices), low pass filtered at either 5 kHz (for neuronal intrinsic property/current-clamp recordings) or 2 kHz (for neuronal and RGC mEPSC recordings), and digitized at 5 kHz (spontaneous neuronal voltage-clamp recordings), 10 kHz (mEPSC recordings) or 50 kHz (intrinsic property current-clamp recordings) through a National Instruments BNC-2090 analogue-digital interface (National Instruments). All RGC and neuronal mEPSC and spontaneous activity experiments were recorded using WinEDR software (V3.7.5), and all neuronal intrinsic property/current-clamp experiments were recorded using WinWCP (V5.2.7; both Strathclyde Electrophysiology Software). Perfusion of aCSF was achieved using a gravity fed set-up, at a rate of ∼2 mL/min, with the inflow positioned ∼2 mm away from the cell and the outflow located ∼10 mm on the opposing side to achieve efficient flow across the recorded cell. The perfusion system had six aCSF holding chambers, which were individually controlled using a six-channel valve controller (Warner Instrument Corporation) to rapidly apply drugs or change recording aCSF solutions. Recording electrodes for whole-cell patch-clamp were pulled on a Model P-87 Flaming / Brown Micropipette Puller (Sutter Instruments), using thick-walled borosilicate glass of dimensions: 1.5 mm OD x 0.86 mm ID (Harvard Apparatus).

To record intrinsic properties neurons were whole-cell voltage-clamped at -60 mV, and their RMP taken immediately upon break-in. From voltage-clamp at -60 mV a current-voltage relationship protocol was run ( -80 to +50 mV, 5 mV steps) before current-clamping the neurons at -60 mV. A current injection stimulus protocol was run to determine the rheobase and to generate the frequency-input relationship. Current was injected for 0.5 s pulses, from -50 pA to +140 pA in 10 pA steps, with 5 seconds recovery at current-clamp I = -60 mV between each injection. The protocol was repeated to take an average result, with the access resistance checked between each recording. For drug application experiments the above procedure was followed, before the RMP and current voltage relationships were re-taken. Standard aCSF perfusion was then switched to the drug containing perfusion, and after 5 minutes of drug application the RMP and current-voltage relationships were repeated. The cell was then returned to current-clamp I = -60 mV and the current injection protocol repeated a further two times. The drugs used in this experiment were tertiapin Q (15 nM, abcam), ML 133 hydrochloride (4 μM, Tocris) and BaCl_2_ (5 μM, Sigma). Cells whose access resistance changed >30% during recording were discarded.

To record spontaneous activity and mEPSCs The resting membrane potential was taken immediately after break-in and neurons were then voltage-clamped at -60 mV. The spontaneous activity was first recorded with standard aCSF perfusing, with a test pulse injected every 2 minutes to monitor the access resistance. Recording times varied from 2-10 minutes depending on the activity level of the cell. For mEPSC recordings the standard aCSF was then switched for aCSF containing 300 nM tetrodotoxin (TTX) and 50 μM picrotoxin (PTX, Tocris), and mEPSCs were recorded for a further 2-10 minutes, checking the access every 2 minutes. If the access resistance was >30 MΩ or changed >20% during recording the cell was discarded.

To measure Kir currents, Neurons were patched in an internal solution that contained (in mM) (K-gluconate 141, NaCl 2.5, HEPES 10, EGTA 11; pH 7.3 with KOH) and an external ACSF described above plus 300 nM TTX, 20 µM NBQX and 50 µM AP5. If membrane potential < K^+^ reversal potential (E_K_), Kir channels conduct in an inward-rectified fashion. To achieve this, neurons were voltage clamped at -60 mV and perfused with an additional 30 mM KCl inducing a calculated E_K_ shift from - 99 mV to -37 mV. A steady-state KCl current was measured, 100 µM BaCl_2_ was perfused for 5-10 seconds in ACSF alone and then remeasured in the presence of KCl to determine the current blocked by Ba^2+^. This block was measured twice and was completely reversed by washout. A similar protocol was implemented for the specific antagonists 150 nM Tertiapin-Q + 20 µM ML-133, except the pre-incubation time was increased to 50 seconds and no washout was attempted.

### Western blotting

Samples were collected in RIPA buffer and stored at -20 °C. On the day of use, the samples were defrosted on ice and a colorimetric BCA assay (Pierce) was run to determine the protein concentration of the samples (measured with a FLUOstar Omega microplate reader). Aliquots of the samples were then taken and diluted in dH2O to a concentration of 32.5 μg/65 μL. Reducing agent (10 μL, NuPAGE) and sample buffer (25 μL, NuPAGE) were added to 65 μL of the diluted sample (final protein concentration 10 μg/20 μL). The samples were then vortexed, spun down and boiled for 10 minutes. Samples were vortexed again before being loaded into a 4-12% Bis-Tris Protein Gel (NuPage), at a volume of 20 μL per well, with ladder loaded into the outside lanes (10 μL, SeeBlue). Electrophoresis was run using MOPS buffer at 120 V. Once the gel had run, proteins were transferred onto a methanol activated PVDF membrane (Millipore) using transfer buffer (96 mM glycine, 12 mM Tris and 20% Methanol) at 80 V for 1 hour. Following transfer, the membrane was blocked in TBS-T solution (20 mM Tris, 137 mM NaCl and 0.1% Tween-20, pH 7.6) containing 5% milk for 1 hour at room temperature. The membrane was then cut in half along the ∼35 kDa level, and each half incubated in its respective primary antibody overnight at 4 °C. Primary antibodies used were: Anti-Calmodulin (1 μg/mL; Merck 05-173) and Anti-GIRK1 (1:5,000; ab129182), Anti-Akt (1:1000, Cell Signaling #9272), Anti-phospho (Ser 473) Akt (1:1000, Cell Signaling #9271).The following day the membranes were washed 3 times in TBS-T, and then incubated for 2 hours at room temperature in 10 mL of TBS-T +5% milk block with the appropriate HRP-linked secondary antibodies. Secondaries used were anti-rabbit IgG HRP-linked antibody (1:1000, Cell Signalling Technologies 7074) and polyclonal goat anti-mouse HRP (1:500, Dako P0447). Following secondary application, the membrane was washed 3 times in TBS-T, exposed to enhanced chemiluminescent reagents (LumiGlo) for 1 minute, and developed manually on Carestream BIOMAX light film. Blots were digitally scanned and densitometric analysis was performed using ImageJ.

### Conditioned media experiments

To generate astrocytic conditioned media (ACM) for application to neuronal cultures, on astrocyte DIV11 the flasks were split and plated down in 1% NBA instead of DMEM. On neuronal DIV0 a proportion of this astrocytic media was pipetted from the astrocytes’ wells, collected in a falcon tube, and dosed with AraC. After aspirating Opti-MEM from the neuronal samples, this treated ACM was used to feed the neurons instead of standard 1 % NBA. Astrocyte plates were topped up with fresh 1% NBA and maintained alongside neurons. For neuronal DIV4 feeding, media was again collected from the astrocyte plate, treated with AraC, and fed to the neurons. To generate ACM for mass spectrometry analysis, on astrocyte DIV7, 1x 75 cm^2^ flask was split and plated into 3x 75 cm^2^ flasks, and on DIV11 these flasks were shaken at 140rpm for 1hour at 37°C and media aspirated before remaining astrocytes being plated down on 6-well plates in NBA 1% media. After 72 hours the NBA 1% media was removed, and replaced with their respective collection medias. The two different sample collection medias used were Neurobasal-A minus phenol red (Gibco) with added glutamine (1 mM) and Anti-Anti (1%), and phenol-red free DMEM (Gibco) with added Anti-Anti (1%). One day later the sample was collected from each plate into chilled 50 mL falcon tubes containing protease inhibitor (cOmplete). All following steps were carried out on ice. Samples were centrifuged at 800 rpm for 4 minutes at 4 °C and then the remaining supernatant centrifuged again at 1000 rpm for 5 minutes at 4 °C, thereafter the supernatant was passed through a sterile 0.22 μm filter (Fisher) into a chilled falcon. Twelve mL of the sample was then added to the reservoir chamber of a pre-chilled 3K centrifugal tube (Amicon), before being centrifuged at 4,000 g for 40 minutes at 4 °C. The filtrate was discarded and the concentrated sample was then ultracentrifuged at 50,000 rpm for 1 hour at 4 °C. The concentrated samples were recovered into chilled Eppendorf tubes, a BCA assay run to determine protein concentration, before being stored at -80 °C. To generate neuronal conditioned media, 12 mL of rat suspension was plated into a 6-well plate and fed with 1% NBA + AraC on DIV0. On DIV3 the sample conditioning media (phenol red-free NBA), was added and on DIV4 the media was collected and concentrated as per the ACM protocol. Where performed protease digestion of ACM used proteinase K-agarose (Sigma) (1 Unit per 5 ml media) for 8 h at 37 °C with constant shaking. The agarose-bound enzyme was then removed by brief centrifugation and 50 µM proteinase K inhibitor (Calbiochem) added to supernatant.

### Immunohistochemistry and confocal imaging

This was carried out as described ^65–67^. Cells were fixed in 1% formaldehyde for 20 minutes, washed 3 times with PBS, permeabilised with 0.5% NP40 (Life Technologies) for 5 minutes and then blocked in 1% BSA (Sigma) for 15 minutes. Antibodies were applied in 1% block solution, with 300 μL of solution applied to each well. The following antibodies were included in the solution: Synapsin 1 polyclonal rabbit antibody (1:1,000, SySy 106 103), Homer 1 monoclonal mouse antibody (1:200, SySy 160 011) and fluorescein isothiocyanate (FITC)-conjugated GFP (1:500, abcam, ab6662). The plate was wrapped in foil and left to rotate at 4 °C overnight. The following morning the primary antibody solution was collected and saved, the wells washed three times with PBS, and then 300 μL of 1% block solution containing secondary antibodies was applied to each well. The secondaries used were: anti-mouse Cy3 (1:500, Jackson ImmunoReseach, 115-165-044) and anti-rabbit Alexa Fluor 647 (1:1,000, abcam, ab150083). The plates were left rotating in secondary antibodies for two hours at room temperature. The coverslips were then washed five times in PBS and mounted on glass microscopy slides using Vectashield mounting media (with DAPI). The slides were covered and sealed from light and left at room temperature for two hours for the mounting medium to set, then stored at 4 °C in a slide box. The samples were later imaged for synaptic markers on a Nikon A1R FLIM confocal microscope. GFP positive cells were selected under 10x magnification, before switching to an oil-submerged 60x lens to record processes of interest. For each recorded cell, 3 dendrites were selected and imaged at approximately 100 μm out from the cell body, which was typically on a secondary or tertiary branch. Around five cells were imaged per coverslip, across three independent culture batches. Images were analysed offline in Fiji (ImageJ), counting co-localised pre- and post- synaptic markers that occurred along the GFP positive dendrite region imaged. Co-localised puncta appeared as purple-yellow marks, where the pre-synaptic (blue) post-synaptic (red) and dendrite (green) markers overlapped. The number of synapses counted for each dendrite was divided by the length of the region and multiplied to give the number of synapses per 10 μm length. The average number of synaptic puncta/10 μm across the three dendrites per cell was taken to get the cell average.

### RNA Extraction

RNA extraction was carried out using the High Pure RNA Tissue kit (Roche) as per manufacturer’s instructions. In brief, samples were homogenised in 700 μL of Lysis re-agent where RNA is extracted and purified by binding and washing through an RNeasy Mini-spin column. Specifically, homogenised samples were centrifuged at 8,000 g for 15 seconds before discarding of filtrate and addition of 100 μL DNAse solution (10μL DNAse I mix and 90μL DNAse incubation buffer) which was then left to incubate at room temperature for 15 minutes. Sequential washes were then performed using 500μL of wash buffer I at 8,000g for 15 seconds, 500μL of wash buffer II at 8,000g for 15 seconds, and 200μL of wash buffer II for 2 minutes at 13,000g. The final purified RNA was eluted in 42 μL elution buffer at 8,000g for 1 minute. Low-yield RNA extraction (for example following ribosomal pulldown) was carried out using the Absolutely RNA Nanoprep kit (Agilent) as per manufacturer’s instructions. Samples were lysed in 100 μL of lysis buffer with β-mercaptoethanol; mixed with equal volume of 80% sulfolane and added to silica-based fibre filter columns. DNAse treatment was carried out for 15 mins; columns were washed and dried, and RNA eluted in 20 μL of pre-warmed 60⁰C RNAse free water.

### Chromatin Immunoprecipitation (ChIP) Seq

Cells were cultured in 15cm plates – astrocyte-neuron coculture, and pure astrocytes and pure neurons monoculture, and were retrieved on DIV10. Chromatin immunoprecipitation was performed with the SimpleChIP® Plus Enzymatic ChIP Kit (Magnetic Beads) (Cell Signalling Technology, #9005) according to the manufacturer’s protocol. For each 15cm culture dish, 40mls of 1X PBS (Life Technologies) were placed on ice. A further 10mls of 1X PBS with 50µl of 200X Protease Inhibitor Complex (PIC) (#7012) was prepared for each 15cm culture dish and place on ice. 540µl of 37% Formaldehyde (Sigma Aldrich) was added to each 15cm culture dish containing 20ml of culture media (final Formaldehyde concentration per dish is 1%). The plates were gently swirled to mix and allowed to incubate at room temperature for 10 minutes. 2mls of 10X Glycine (#7005) was then added to each dish and the plates were gently swirled to mix. The plates were left to incubate at room temperature for 5 minutes. The media was removed, and cells were washed with 20mls of ice-cold 1X PBS. The plates were gently swirled, and PBS aspirated. The wash step was repeated once more, and PBS aspirated. 5mls of PBS with PIC was then transferred into each plate and using a cell scraper, the cells were transferred into a 15cm conical tube. The tubes were centrifuged at 2000*xg* for 5 minutes at 4°C.

For each immunoprecipitation reaction (IP), we prepared 3 buffers: A (250µl of 4X Buffer A (#7006), 0.5µl 1M dithiothreitol (DTT) and 5µl of 200X PIC were added to 750µl of water); B (275µl of 4X Buffer B (#7007) and 0.55ml of 1M DTT were added to 825µl of water) and ChIP buffer (10µl of 10X ChIP Buffer (#7008 and 0.5µl 200X PIC were added to 90µl of water). The supernatant was removed, and cells resuspended in 1ml of ice-cold 1X Buffer A per IP prep. The samples were incubated on ice for 10 minutes and mixed by inverting the tubes every 3 minutes. The nuclei were then pelleted by centrifugation at 2000*xg* for 5 minutes at 4°C, and supernatant removed. The pellet was resuspended in 1ml of ice-cold 1X Buffer B per IP prep. The tubes were centrifuged at 2000*xg* for 5 minutes at 4°C, and supernatant removed. The pellet was resuspended in 100µl of 1X Buffer B per IP prep. 1.0µl of Micrococcal Nuclease (#10011) was added to each IP prep, and mixed by inverting the tube several times. The tubes were incubated in a heating cabinet at 37°C for 20 minutes. 10µl of 0.5M EDTA (#7011) was added to each tube to halt the reaction. The tubes were immediately placed on ice for 2 minutes. The nuclei were then pelleted by centrifugation at 16,000*xg* for 1 minute at 4°C and supernatant removed. The pellet was resuspended in 100µl of 1X ChIP Buffer per IP prep and incubated on ice for 10 minutes. Lysates were sonicated with the Bioruptor® Sonication System (Diagenode), set at HIGH setting for 3 cycles of 30 seconds ON followed by 30 seconds OFF. Lysates were clarified by centrifugation at 9400*xg* for 10 minutes at 4°C. The supernatant, which is the cross-linked chromatin preparation, was transferred into a fresh Eppendorf tube. 25µl of the chromatin preparation was removed for Analysis of Chromatin Digestion and Concentration. The rest of the chromatin preparation was stored at -80°C until ready for use. The following were added to 25µl aliquot of chromatin preparation – 3µl 5M NaCl (#7010) and 1µl RNAse A (#7013). The mixture was vortexed to mix and incubated at 37°C for 30 minutes. 1µl of Proteinase K (#10013) was added to each tube, vortexed to mix and incubated overnight at 65°C. DNA was purified and resolved on a 1.3% agarose gel with a 100bp DNA marker to check fragmentation (target 150-900bp). DNA concentration was determined using the Qubit™ dsDNA HS Assay Kit (Thermo Fisher Scientific) and Qubit™ Fluorometer (Thermo Fisher Scientific).

For each IP 100µl (5-10µg) of digested, cross-linked chromatin preparation was added to 400µl of 1X ChIP Buffer. 10µl of the diluted chromatin (2%) was transferred into a fresh Eppendorf tube labelled “Input” and stored at -20°C until ready for use. 500µl of the diluted chromatin was transferred to a 1.5ml microcentrifuge tube and the antibody against the histone marks of interest were added (Table 3). The samples were incubated overnight on a rotating wheel at 4°C (Positive Control Histone H3 (D2B12) Rabbit mAb (#4620), Negative Control Normal Rabbit IgG (#2829), H3K27me3 (Abcam ab6002), H3K27ac (Abcam ab4729)) ChIP-Grade Protein G Magnetic Beads (#9006) were gently vortexed to resuspend the beads. 30µl of the magnetic beads were then immediately added to each IP reaction and incubated for 2 hours on a rotating wheel at 4°C. The washing steps were performed in the cold room (4°C). Firstly. the beads were washed with 1ml of Low Salt Wash and incubated for 5 minutes on a rotating wheel. The tubes were briefly centrifuged (3 seconds at 2000*xg*) and recovered on a magnetic rack. The supernatant was discarded, and beads resuspended in a further 1ml of Low Salt Wash. The wash step was repeated to complete a total of 3 washes with Low Salt Wash and 1 wash with High Salt Wash.

After the final wash with High Salt Wash, the beads were resuspended in 150µl of 1X ChIP Elution Buffer. The tubes were placed on the thermomixer (Eppendorf Thermomixer C) at 400rpm and 65°C for 30 minutes to elute the chromatin from the magnetic beads. The magnetic beads were pelleted on a magnetic separation rack. The eluted chromatin was carefully transferred into fresh Eppendorf tubes. 150µl of 1X ChIP Elution Buffer was also added to the 2% Input tubes and were set aside at room temperature until ready for use. 6µl of 5M NaCl and 2µl of Proteinase K were added to all tubes (eluted chromatin and 2% input) and incubated overnight at 65°C to reverse the cross-link. The samples were purified using the DNA spin columns provided in the kit.

DNA was purified using the spin columns supplied in the kit, according to manufacturer’s protocol. 375µl of DNA Binding Buffer (#10007) was added to each sample and vortexed briefly. 450µl of each sample was transferred to a DNA spin column/collection tube assembly. The tubes were centrifuged at 18,500*xg* for 30 seconds at room temperature. The flow-through was discarded and this step was repeated with the remaining sample. 750µl of Wash Buffer (#10008) was added to the spin column. The tubes were centrifuged at 18,500*xg* for 30 seconds and flow-through discarded. The tubes were centrifuged at 18,500*xg* for a further 30 seconds to dry out the membrane. The collection tube was discarded, and the spin column was placed in a clean Eppendorf tube. 30µl of DNA Elution Buffer (#10009) was added to each spin column. The tubes were centrifuged at 18,500*xg* for 30 seconds to elute the DNA. The eluted DNA was passed through the column again to increase yield.

Library preparation for the samples was performed using NEBNext Ultra II DNA Library Prep Kit for Illumina (New England Biolabs Inc, #E7645S) according to the manufacturer’s protocol. 50ng of ChIP-DNA was end-repaired, 5’ phosphorylated and 3’dA-tailed. NEBNext hairpin adaptors containing uracil were ligated to the dA-tailed fragments before excision of the U nucleotide in the adapters. Adapter-ligated libraries were then purified and size-selected to enrich for 150bp adapter-ligated fragments with AMPure XP Beads. Adapter-ligated DNA was then amplified for 7 cycles with indexed i7 and universal i5 primers. Following amplification, libraries were then purified using AMPure XP beads. Libraries were quantified by fluorometry using Qubit dsDNA HS assay and assessed for quality and fragment size using the Agilent Bioanalyser with the DNA HS Kit (#5067-4626) ChIP-sequencing was performed using the NextSeq 500/550 High-Output v2.5 (75 cycle) Kit (#20024906) on the NextSeq 550 platform (Illumina Inc, #SY-415-1002). Libraries were combined in three equimolar pools based on the library quantification results and run across three High-Output Flow Cells.

### RNA-seq and ChIP-seq plus read analysis

The approach is described extensively elsewhere ^15,68^. Briefly, pure rat neuronal cultures, or cultures of rat neurons on mouse astrocytes, were prepared as described above. The RNA was extracted from the samples, converted into complementary strand DNA (cDNA), and sequenced by Edinburgh Genomics. The bioinformatic analysis on the returned sequencing was conducted by ourselves. At-least 1 μg RNA per sample was utilised, with RNA-integrity number (RIN) > 7. RNA-seq reads were mapped to genome sequences using STAR (Spliced Transcripts Alignment to a Reference, ^69^). Per-gene read counts were summarised using featureCounts ^70^, and differential expression analysis performed using DESeq2 ^71^, with a significance threshold calculated at a Benjamini–Hochberg-adjusted P value of <0.05. Gene ontology enrichment analysis was performed using topGO ^72^. For ChIP-seq, reads were mapped to genome sequences using Bowtie2 ^73^, and mixed-species sorting was again carried out using Sargasso. Peaks of sequencing reads, indicating bound histone marks, were called using MACS2 ^74^. Computation of differentially bound sites between conditions was performed with the DiffBind R package ^75^. Differentially bound sites were then annotated using the ChIPSeeker R package ^76^. Mixed-species sorting of RNA-seq or ChIP-seq reads was carried out using the Sargasso (Sargasso Assigns Reads to Genomes According to Species-Specific Origin) Python tool as described ^15^ (source code available: http://statbio.github.io/Sargasso), which utilises a strategy that aims to minimize reads misallocated to the incorrect species whilst maximizing the number of reads unambiguously assigned to the correct species.

### Quantitative RT-PCR

cDNA was generated using the Transcriptor First Strand cDNA Synthesis Kit (Roche). 7 μL of RNA was added to the RT and buffer mixture prepared with random hexamers and oligoDT primers as per kit instructions, and rtPCR carried out with the following programme: 10 min at 25°C, 30 min at 55°C and 5 min at 85°C. qPCRs were performed on a Mx3000P QPCR machine (Agilent Technologies) using the FastStart Universal SYBR Green QPCR Master (Rox) (Roche) reagent. 6 ng of cDNA was used for each reaction and all qPCR results were carried out in duplicate or triplicate, along with no template controls and no RT controls where appropriate. The following cycling programme was used: 10 min at 95°C; 40 cycles of: 30 s at 95°C, 40 s at 60°C (with fluorescence detection), 1 min at 72°C; ending with dissociation curve: 1 min at 95°C and 30 s at 55°C with a ramp up to 30 s at 95°C with fluorescence detection. All data was normalised to house-keeping gene controls (Rpl13a). Primer sequences are detailed below: rKCNJ3 Forward: 5’-TCGTGTTTGAATCTGGATCTC-3’ rKCNJ3 Reverse: 5’-GCCGTGCTGCACATT-3’. rKCNJ4 Forward: 5’-AGATTGATGAGGACAGTCCAC-3’. rKCNJ4 Reverse: 5’-CATCTTGCTCTCTTGAAGCTC-3’. rRPL13A Forward: 5’-CGCACAAGACCAAAAGAG-3’. rRPL13A Reverse: 5’-GTTTCCTTAGCCTCAAGAGC-3’.

### Reverse Phase Protein Array (RPPA)

RPPA of rat cortical neuronal protein lysate was performed as described ^77^. Briefly, 1% Neurobasal A was conditioned by mouse cortical astrocytes for 72 h, and added to rat cortical neurons. After 15 minutes incubation, neurons were lysed in RIPA buffer (50 mM Tris-HCl, 150 mM NaCl, 1mM EDTA, 1% NP40, 1% Na-deoxycholate, 0.1% SDS, pH 7.4) with fresh protease and phosphatase inhibitors added (Roche). Lysates were scraped off the plate, centrifuged at 13,000 rpm for 10 minutes at 4 °C, with insoluble pellet discarded. Protein concentration was determined using Pierce BCA assay (Thermo), adjusted to 1.6 mg/ml with lysis buffer and mixed with 4XSDS sample buffer. Protein was denatured at 95 °C for 5 min and printed as a 4 dilution concentration series onto ONCYTE SuperNOVA nitrocellulose slides at 500 µm spot-to-spot distance. Sample printing was repeated to give 3 technical replicates, with 4 biological replicates per sample. 120 antibodies were profiled using standard procedures, 1 hour incubation with primary followed by 30 min incubation with Dylight-800 anti-species secondary antibodies, with arrays imaged using an Innopsys Innoscan 710 scanner. Sample loading on arrays was determined by staining one slide with fast green protein dye. Images were analysed using Mapix software (Innopsys), to determine fluorescence intensity of each deposition. R2 values of dilution series were recorded and technical replicates demonstrating R2 <0.8 were excluded from further analysis. Median values of the 4-point dilution series were calculated to give a single measure of fluorescence intensity and normalised to fast green total protein stain.

### Calcium imaging

Ca^2+^ imaging was performed as described ^78,79^ at 37°C in aCSF (150 mM NaCl, 3 mM KCl, 10 mM HEPES, 2 mM CaCl2, 1mM MgCl2, 1mM glucose). Briefly, cells were loaded with 11µM Fluo-3 AM (from a stock solution of 2.2 mM Fluo-3 dissolved in anhydrous DMSO containing 20% (w/v) Pluronic detergent) for 30 min at 37°C. Fluo-3 fluorescence images (excitation 472 ± 15 nm, emission 520 ± 15 nm) were taken at one frame per 5 sec using a Leica AF6000 LX imaging system, with a DFC350 FX digital camera. To calibrate images, Fluo-3 was saturated by adding 50 μM ionomycin to the perfusion chamber (to obtain Fmax) and quenched with 10 mM MnCl2 + 50 μM ionomycin to levels corresponding to 100 nM Ca^2+ 80^, which was in turn used to calculate Fmin. Free Ca2+ concentrations were calculated from fluorescence signal (F) according to the equation [Ca^2+^] = Kd(F – Fmin)/(Fmax – F), and expressed as a multiple of the Kd of Fluo-3 (which is approximately 315 nM).

### Data analysis and statistics

Electrophysiology analysis was run using Stimfit (v0.15.4) software. In-built analysis functions were used to extract the data from IV relationships and FI protocols, which was then exported to Excel for further analysis. For spontaneous activity and mEPSC analysis, a custom noise filtering script was written and implemented in Stimfit. Basically, a Fast Fourier Transform is run over the recording to detect constant sources of noise. The noise waves are then subtracted from the recording, to improve the clarity of the trace. For spontaneous activity, large events that have over 200 pC charge passing through them are first detected, removed and recorded as events, providing details such as the charge and amplitude of each event. A mini analysis script is then run to detect remaining small spontaneous events, extracting them and recording details including their amplitude, time constant and charge. The total charge, from both large and small events, for the recording is than calculated, as well as the average amplitude from all events. For mEPSC recordings, the noise protocol and mini analysis script is run. The results are then exported to Excel for further analysis. All results are given as mean ± standard error of mean (SEM), unless otherwise stated. All statistical tests employed are stated in the figure legends.

### Data availability

RNA-seq and ChiP-seq data in this study have been deposited at the EMBL European Bioinformatics Institute Biostudies platform under accession number E-MTAB-14070 (RNA-seq) and E-MTAB-14081 (ChIP-seq).

---

**Figure S1.**
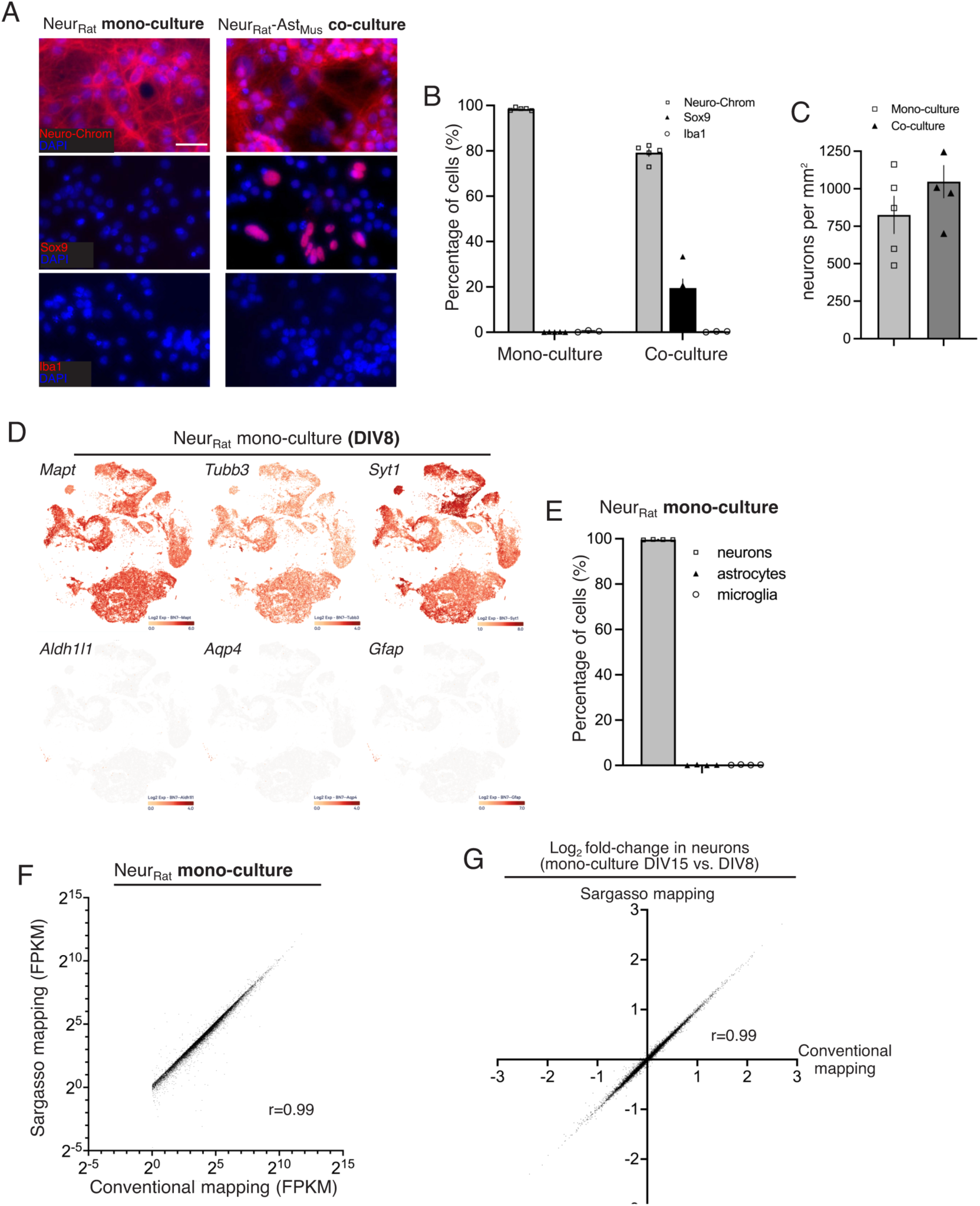
Related to Fig. 1. **A, B).** Characterisation of the proportion of neurons, astrocytes and microglia in neuronal mono-cultures and neuron-astrocyte co-cultures. Scale bar: 50 µm. **C)** Density of Neuro-Chrom positive neurons in neuronal mono-cultures and neuron-astrocyte co-cultures. **D, E)** Single nucleus RNA-seq of neuronal mono-cultures was performed and the proportion of neurons, astrocytes and microglia quantified. (D) shows examples of neuron- and astrocyte-specific gene expression. **F)** An overview of the impact of the species-specific Sargasso workflow on reported FPKM values. FPKM values of genes in rat neuron mono-cultures when mapped by conventional STAR mapping vs. Sargasso mapping which requires rat-neuron read disambiguation prior to STAR mapping. **G)** An overview of the impact of the species-specific Sargasso workflow on reported fold- changes in differential gene expression reporting. Differential gene expression in rat neuronal mono-cultures (DIV15 vs. DIV8) was calculated using the Sargasso workflow and also by conventional mapping, prior to DESeq2 differential gene expression analysis, and compared.

**Figure. S2.**
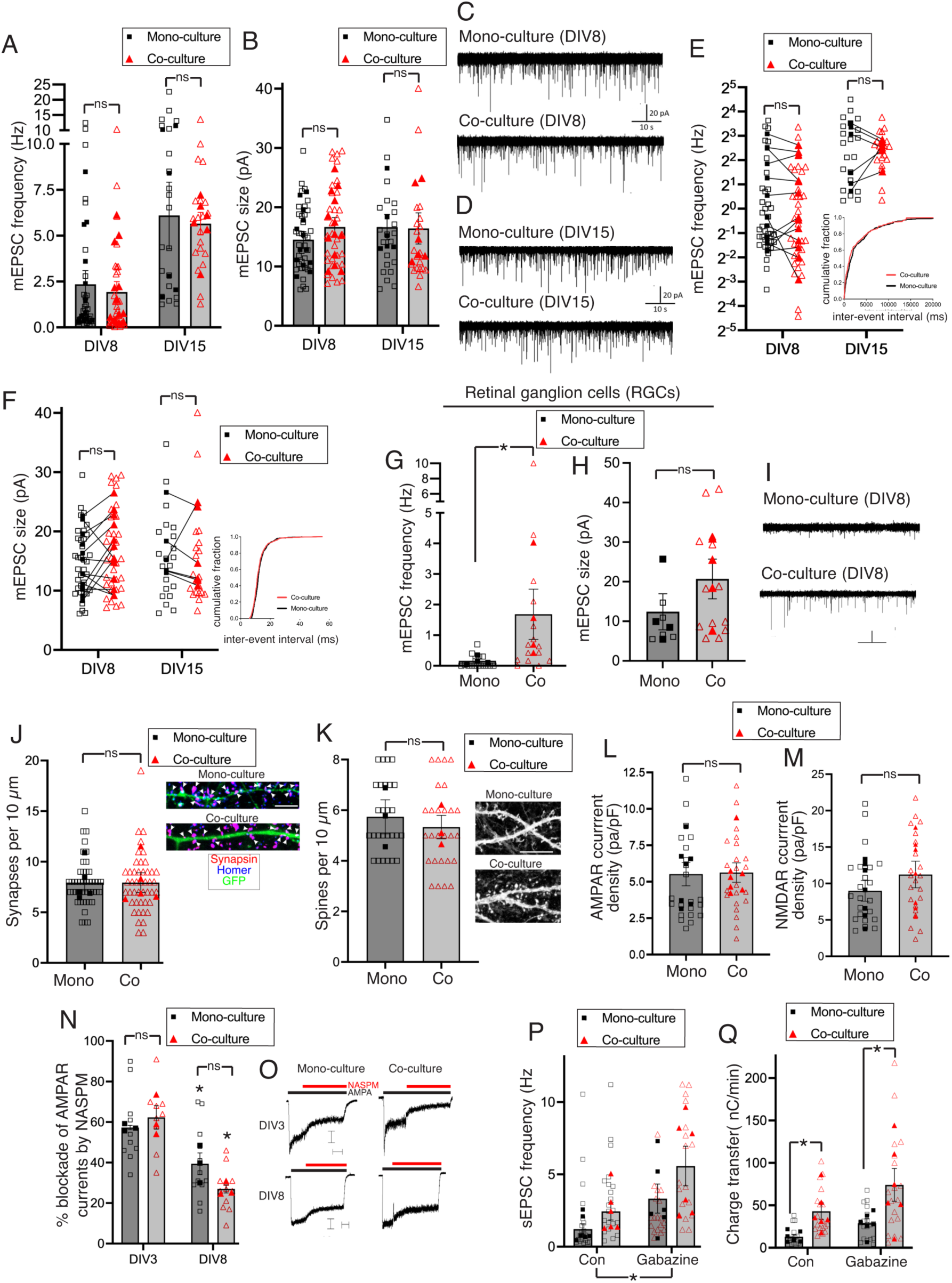
Related to Fig. 3. **A, B)** Miniature EPSC frequency (A) and amplitude (B) was recorded at the indicated age in the presence or absence of astrocytes. To test the effect of co-culture we used two approaches: a hierarchical nested t-test (treating cells within the same independent culture as technical replicates) and a two-way ANOVA on the mean value for each independent culture (a third paired approach is shown in Fig. S2a). DIV8: t=0.404, df=24, p=0.690. DIV15: t=0.44, df=10, p=0.669. Astrocyte effect: F (1, 17) = 0.6428, p=0.434; age effect: F (1, 17) = 9.426, p=0.007, 2-way ANOVA. B) DIV8: t=0.9847, df=24, p=0.335. DIV15: t=0.1535, df=10, p=0.881. Astrocyte effect: F (1, 34) = 0.2513, p=0.619; age effect: F (1, 34) = 0.2409, p=0.683, 2-way ANOVA. N=13 (DIV8) and 6 (DIV15) independent cultures. Individual cell values are shown as open symbols, with the mean of each culture as filled symbols, here and throughout. **C, D)** Example recordings relating to (A) and (B). **E)** Relating to the data presented in Fig. S2a, a paired analysis was performed on mono- vs. co-culture mEPSC frequency, pairing data obtained from the same neuronal culture. DIV8: t=1.265, df=12, p=0.230 (n=13); DIV15: t=0.3054, df=5, p=0.772 (n=6), paired t-test. Inset shows an example cumulative distribution curve. **F)** Relating to the data presented in Fig. S2b, a paired analysis was performed on mono- vs. co-culture mEPSC frequency, pairing data obtained from the same neuronal culture. DIV8: t=1.640, df=12, p=0.127 (n=13); DIV15: t=0.1168, df=5, p=0.912 (n=6), paired t-test. Inset shows an example cumulative distribution curve. **G, H)** Effect of astrocytes on synaptic properties in retinal ganglion cells: mEPSC frequency (G) and amplitude (H). K: P=0.029, Mann Whitney test, sum of ranks: 10 (mono-culture) and 26 (co-culture, n=4 cultures, 3-5 cells per culture), n=4 cultures. L: t=1.245, df=6, p=0.260, n=4 cultures (same cells analysed as in (G) but a reduced number of data points is due to some cells not exhibiting any mEPSCs). **I)** Example traces relating to (G) and (H). Scale bars: 50 pA, 5 s. **J)** Neurons were sparsely transfected with eGFP and stained for the indicated pre- and post-synaptic markers, with co-localized puncta along the shaft scored as a functional synapse. Nested t-test: t=0.4101, df=8, p=0.693 (10 cells per culture per condition analysed, n=5). Scale bar: 5 µm. **K)** Neurons were sparsely transfected with eGFP and spines counted. Nested t-test: t=0.4985, df=4, p=0.644 (9 cells per culture per condition analysed, n=3). Scale bar: 10 µm**. L, M)** 50 µM (S)-AMPA and 150 µM NMDA receptor whole-cell currents were recorded at DIV8 cultures ± astrocytes. Nested t-test: t=0.08142, df=12, p=0.937 (G); t=0.9074, df=12, p=0.382 (H), n=7. **N)** Effect of NASPM (1-Naphthylacetyl spermine) on AMPA receptor-mediated currents. Age effect: F (1, 8) = 41.73, p=0.0002 2-way ANOVA followed by Sidak’s post-hoc test, *p=0.030, 0.0006 (left to right). Astrocyte effect: F (1, 8) = 0.8112, p=0.394. Nested t-test: t=0.5303, df=4, 0.624 (DIV3); t=2.110, df=4, p=0.103. **O)** Example traces. Scale bars: 25 pA (upper), 100 pA (lower), 3 s (both). **P, Q)** Effect of gabazine on spontaneous EPSC frequency (P) and EPSC charge transfer (Q) in the presence or absence of astrocytes. P: Gabazine effect: F (1, 10) = 5.479, p=0.041. Q: astrocyte effect: F (1, 10) = 19.20, p=0.001, *p=0.033, 0.008 (Holm Sidak’s post-hoc test). N=6 for all conditions/treatments.

**Figure S3.**
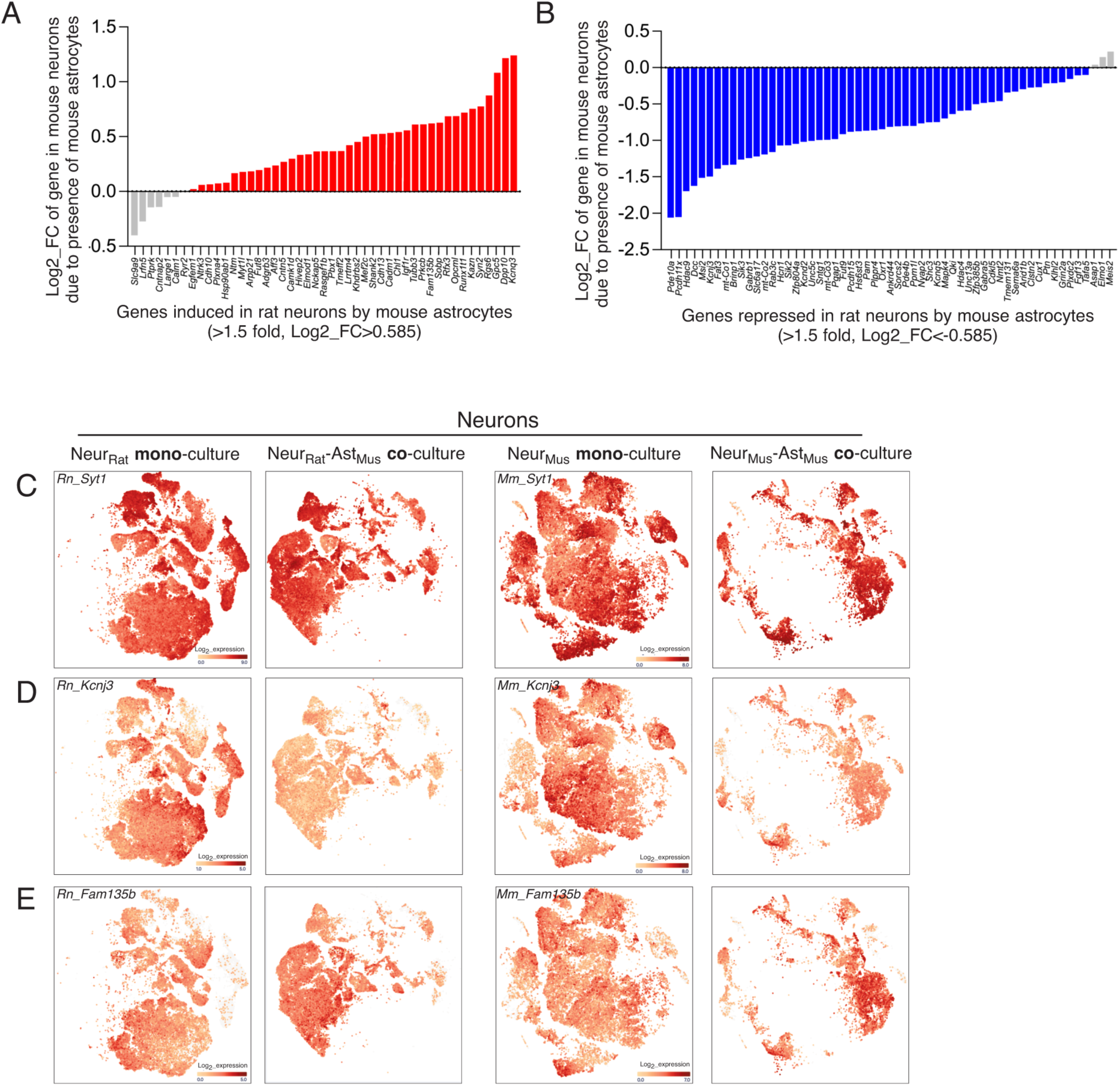
related to Fig. 6. **A, B)** Single nucleus RNA-seq was performed on rat and mouse cortical neurons cultured in the presence or absence of mouse astrocytes. Genes induced (A) and repressed (B) > 1.5-fold by mouse astrocytes in rat neurons were taken and the change observed in mouse neurons due to the presence of mouse astrocytes was plotted. As can be seen, up-regulated genes are overall up-regulated in both species, while up-regulated genes are similarly conserved across species. **C-E)** Example sc-RNA-seq representations of the expression of genes in rat (left two columns) and mouse (right two columns) neurons in the presence or absence of mouse astrocytes. A neuronal marker (*Syt1*) is shown in (C), down-regulated gene *Kcnj3* in (D), and up-regulated gene (*Fam135b*) in (E).

